# Enhanced prediction of cognitive function using aging-sensitive networks within the human structural connectome

**DOI:** 10.1101/2019.12.13.875559

**Authors:** James W. Madole, Stuart J. Ritchie, Simon R. Cox, Colin R. Buchanan, Maria Valdés Hernández, Susana Muñoz Maniega, Joanna M. Wardlaw, Mat A. Harris, Mark E. Bastin, Ian J. Deary, Elliot M. Tucker-Drob

**Affiliations:** Department of Psychology, University of Texas at Austin, Austin, TX; Social, Genetic and Developmental Psychiatry Centre, King’s College London, London, UK; Lothian Birth Cohorts, University of Edinburgh, Edinburgh, UK; Department of Psychology, University of Edinburgh, Edinburgh, UK; Scottish Imaging Network, A Platform for Scientific Excellence (SINAPSE) Collaboration, Edinburgh, UK; Centre for Clinical Brain Sciences, University of Edinburgh, Edinburgh, UK; Division of Psychiatry, University of Edinburgh, Edinburgh, UK; Population Research Center, University of Texas at Austin, Austin, TX

## Abstract

Using raw structural and diffusion brain MRI data from the UK Biobank (UKB; *N* = 3,155, ages 45-75 years) and the Lothian Birth Cohort 1936 (LBC1936; *N* = 534, all age 73 years), we examine aging of regional grey matter volumes (*nodes*) and white matter structural connectivity (*edges*) within networks-of-interest in the human brain connectome. In UKB, the magnitude of age-differences in individual node volumes and edge weights corresponds closely with their loadings on their respective principal components of connectome-wide integrity (|*r*_nodes_| = 0.459; |*r*_edges_| = 0.595). In LBC1936, connectome-wide and subnetwork-specific composite indices of node integrity were predictive of processing speed, visuospatial ability, and memory, whereas composite indices of edge integrity were associated specifically with processing speed. Childhood IQ was associated with greater node integrity at age 73. However, node and edge integrity remained associated with age 73 cognitive function after controlling for childhood IQ. Adult connectome integrity is therefore both a marker of early-life cognitive function and a substrate of late-life cognitive aging.

---

Many cognitive abilities exhibit declines across adulthood^1, 2^. These declines have consequences both for individuals, who may be less able to perform important everyday functions^3, 4^, and for aging societies, whose workforce productivity and social and medical resources may be prematurely exhausted^5^. Delineating the neurodegenerative processes underlying aging-related cognitive decline may crucially advance our ability to detect, and ultimately mitigate, prevent, or ameliorate aging-related cognitive impairments.

The human brain exhibits widespread structural changes with aging^6^, the patterning of which is only partly documented. It is not yet known which aging-related changes in brain structure are particularly relevant for adult cognitive functioning. Here, we take a cross-cohort magnetic resonance imaging (MRI) approach to identify elements of brain morphometry and inter-regional white matter connectivity that show sensitivity to aging and are relevant to late-life cognitive functioning. Following recent advances in network neuroscience, we model each participant’s brain as a macroscale *connectome*: a network of discrete grey matter regions (*nodes*) that are connected by bundles of myelinated white matter fibers (*edges*)^7^. In the UK Biobank (UKB) sample, we identify major dimensions of connectome-wide edge and node integrity, which we examine in relation to cross-sectional age trends in connectome elements. Using regression weights obtained from these UKB analyses, we create indices of general dimensions of edge and node integrity in the whole-brain connectome and ten of its subnetworks-of-interest in the narrow-aged Lothian Birth Cohort 1936 (LBC1936; all age 73 years). We use these weighted indices of connectomic integrity to predict core cognitive abilities known to decline with adult age^8,9,10^: processing speed, visuospatial ability, and memory.

The current work extends beyond well-replicated findings that coarse measures of brain structure have moderate associations with age and cognitive abilities. Measures of whole- and regional-brain volumes^11,12,13^ and white matter microstructure^14,15,16^ across multiple major tracts have been linked to cognitive function and age-related cognitive decline. For example, in over 18,000 adult participants in middle and older age, Cox et al.^17^ reported *r* = 0.276 for the relation between total brain volume (TBV) and general cognitive ability. The same study reported that when multiple global indices of grey- and white-matter macrostructure and microstructure were used to predict general cognitive ability in a subsample of older adults, the prediction increased to *r*_multiple_ = 0.361. Moderate to strong associations have also been found between gross MRI measures and age (*r*’s = −0.573 to −0.254)^18^. Taken together, these findings suggest roles for omnibus measures of brain structure in cognitive aging. The relatively low resolution of these indices, however, has constrained the level of specific insight able to be gleaned regarding the neuroanatomical networks relevant for cognitive aging.

Here, we further investigation the neurobiology of cognitive aging by examining the individual elements of the human structural connectome in relation to adult age and late-life cognitive function^19^. We move beyond previous studies, which have largely documented age trends in summary indices of connectome topology (e.g., *strength*, *global efficiency*)^20, 21^, or have used large-scale, exploratory methods to examine how a range of morphometric and diffusion tensor measures relate to age and a broad array of sociodemographic variables^22, 23^. Building on research detailing intrinsic networks within the human *functional* connectome^24, 25^, we create *structural* connectome masks to partition the whole-brain connectome into ten prespecified networks-of-interest (NOIs). Several of these networks have been previously implicated in cognitive function (e.g., Central Executive^26, 27^; Parieto-Frontal Integration Theory (PFIT)^28, 29^; Multiple Demand^30, 31^), whereas others underscore more basic functions (e.g., Salience^32, 33^; Sensorimotor^34, 35^), and therefore serve as negative controls. These subnetworks are distributed throughout the brain and partially overlap. We examine age trends for individual elements within the whole-brain connectome and within each NOI, before exploring how these age trends relate to general dimensions of regional volume and interregional connectivity, respectively. We use summary indices of volumetric structure and white matter connectivity at age 73 years to predict concurrent measures of processing speed, visuospatial ability, and memory, and examine the robustness of these associations relative to controls for TBV and age-11 cognitive function.

## Results

### Description of Structural Connectome Construction and Analyses

Whole-brain structural connectomes were created for each participant in UK Biobank (UKB; *N* = 3,155; ages 45-75 years) and the Lothian Birth Cohort 1936 (LBC1936; *N* = 534; all age 73) from 85×85 matrices of fractional anisotropy weights (*edges*), reflecting strength of connections between cortical and subcortical regional volumes (*nodes*) parceled per the Desikan-Killiany atlas^36^. Masks were created to partition whole-brain connectomes into nine prespecified NOIs (Fig. 1; Table S1; Table S2), including a null network consisting of edges and nodes not contained in any other NOI. Several NOIs were composed of partially overlapping *edges* and *nodes*, collectively referred to here as *elements* (Table S3). Where applicable, results provide details for how overlapping elements were handled.

**Figure 1.**
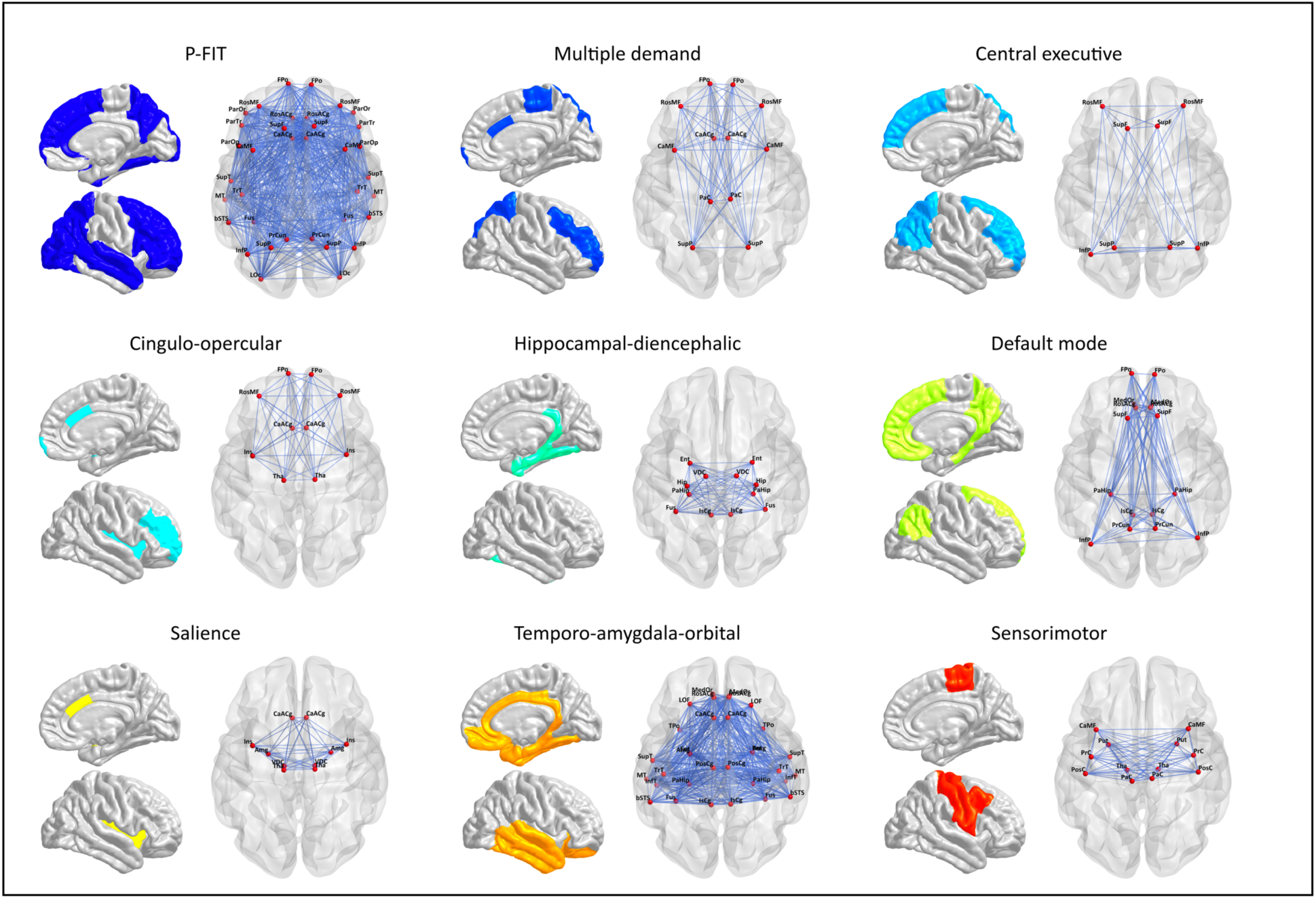
**Anatomical maps of each NOI.** Anatomical maps of each prespecified brain NOI displaying the network-specific connectome elements (*i.e.*, edges and nodes). We also considered a whole-brain network and a null network comprised of elements not belonging to any prespecified network.

Analyses were run using unthresholded matrices, which were determined to be largely similar to consistency-based thresholded^37^ matrices (Fig. S1; Supplementary Method and Results). We performed two sets of analyses, one within-sample and one out-of-sample. Within the large, age-heterogeneous UKB sample, we documented age differences in the volumetric and structural connectivity properties of each NOI. We then assessed whether general dimensions of overall network integrity (Fig. S2) were preferentially associated with age-sensitive elements. In the age-homogeneous LBC1936, we used UKB-trained models of *connectome age* and *connectome integrity* to predict variation in cognitive functions. Finally, we ran a series of sensitivity analyses.

### Connectome Aging

Cross-sectional age-trends in all individual elements were estimated in the UKB structural MRI sample. Density distributions of the element-wise age associations for the whole-brain connectome and each individual NOI are presented in Fig. 2. The majority of elements showed small to modest negative associations with age (*edges*: 2299/3570 [64.4%] < 0, mean *r* = −0.034, range = −0.414 to 0.265; *nodes*: 83/85 [97.6%] < 0, mean *r* = −0.172, range = −0.313 to 0.085). Several subnetworks displayed bimodal distributions of age-node associations, potentially indicating multiple aging-related processes within these networks. Elements within the Central Executive network, a subset of the larger PFIT network, displayed the steepest average age-related gradients (mean *r*_age-edge_ *=* −0.161; mean *r*_age-node_ = −0.242; Table S4), indicating that it demarcates a particularly age-sensitive constellation of elements. Only the Salience network contained a majority of edges with positive age associations (37/45 [82.2%] *r’*s > 0). In contrast, all ten of its nodes displayed negative age associations.

**Figure 2.**
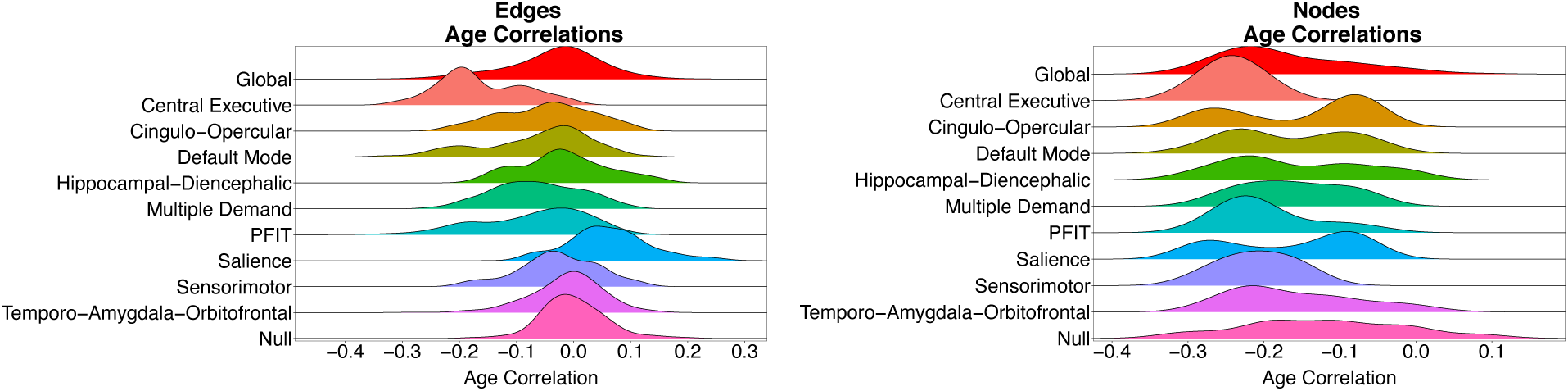
**Density distributions of age associations.** Density distributions of each element’s association with age, categorized by prespecified NOIs. All subnetworks are subsets of the whole-brain (Global) network, such that comparison with the red distribution at the top of both panels is not a comparison of independent elements, but a comparison of a subset to a whole.

#### General dimensions of connectome integrity

The widespread patterns of age-related decrements across NOIs suggests that individual elements may represent broader dimensions of interindividual variation in global connectome integrity. We examined this possibility in edges and nodes separately using PCA (Fig. S2). The first PC accounted for 10.8% and 35.7% of variation in edges and nodes, respectively. The second PC accounted for less than 1/5 the variance accounted for by the first corresponding first Eigen value (Fig. S3). When PCAs were performed on covariance matrices of network-specific composite indices of integrity (Fig. S2d), the first PC accounted for 59.7% and 83.5% of the variation for edges and nodes, respectively (Fig. S4; Tables S5 & S6; Supplementary Results).

Whole-brain loadings were overwhelmingly positive (*edges*: 95.6% of loadings > 0, *nodes*: 100% of loadings > 0). There was considerable heterogeneity in whole-brain loadings across elements within NOIs (Fig. 3). Elements within the Central Executive network displayed the largest average loadings, again suggesting that this small subset of the PFIT network may disproportionately index overall brain integrity. As with the age associations, several networks displayed bimodal distributions of PC loadings, potentially suggesting that the same clusters of elements that show concurrent age-sensitivity also represent equivalent indices of overall network integrity.

**Figure 3.**
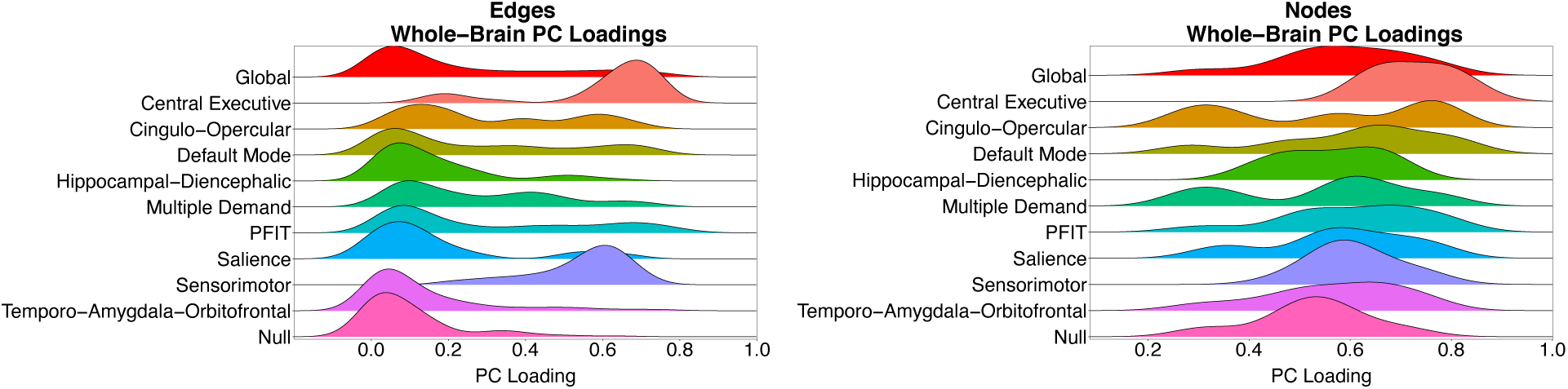
**Density distributions of whole-brain principal component loadings.** Density distributions of loadings on the first principal component of the whole-brain connectome, categorized by prespecified NOIs.

#### Connectome aging occurs along general dimensions of variation in edge and node integrity

We tested the extent to which aging-related differences in individual connectome elements occurred along the general dimensions of edge and node integrity identified above. In UKB, we estimated the correlation between two vectors: (1) a vector of the loadings of each element on the first PC (both whole-brain and network-specific) and (2) a vector of the age correlations between each element and age. We conducted this analysis separately for edges and nodes. Note that because the connectome elements were partialled for age prior to conducting the Eigen decompositions, the resulting association between age-sensitivity and PC loadings is *not* an artifact of age effects driving connectome element covariation.

Fig. 4 displays the association between the loadings and the age correlations for edges (left) and nodes (right) in the whole brain. Edges that had stronger loadings evinced steeper age-gradients (*r* = −0.595): the more indicative an edge was of global variation in brain connectivity, the greater its negative association with age. The same pattern was evident for the nodes (*r* = −0.459): the more representative a node was of global variation in brain volume, the stronger its negative association with age. Similar patterns were obtained when analyses were conducted separately for each individual NOI (Figs. S5 & S6; Supplementary Results).

**Figure 4.**
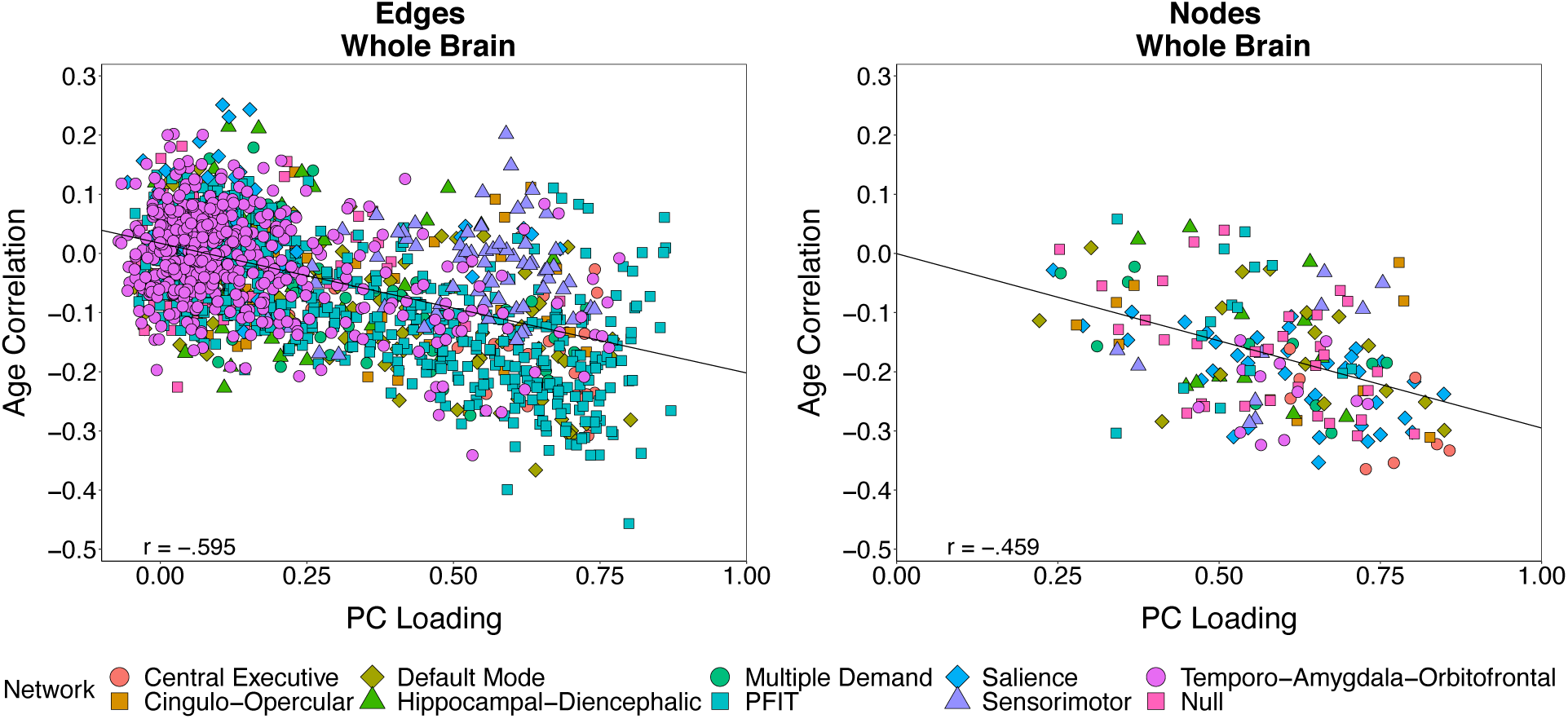
**Scatterplot of age correlations and principal component loadings.** Scatterplots of each connectome element’s correlation with age against its loading on a single principal component (based on an age-partialled correlation matrix (Fig. S2)). Analyses were conducted separately for edges (left) and nodes (right). Each point represents a single element of the connectome (3,564 non-zero edges; 85 nodes). Points are categorized by the NOI to which the element belongs. Elements belonging to multiple NOIs are plotted once for each group membership and jittered for the sake of visual interpretation. Reported correlations and displayed regression lines reflect analyses including each element only once.

We tested whether the observed associations between PC loadings and age correlations were explained by how central the elements were within the whole-brain connectome, a potential metabolic cost that could confer susceptibility to degeneration with age^38^ (Figs. S7 & S8; Supplementary Results). We found that the topological centrality of elements was strongly correlated with loadings on their respective PCs (*r*_edges_ = 0.650; *r*_nodes_ = 0.558; all *p*’s < 0.0005; Fig. S7), but only modestly associated with its age correlation (*r*_edges_ = −0.194, *p* < 0.0005; *r*_nodes_ = −0.235, *p* = 0.031; Fig. S8). Topological connectedness of connectome elements was therefore insufficient to explain associations between PC loadings and age correlations.

#### General Dimensions of Connectome Integrity are Associated with Late-Life Cognitive Function

The finding that connectome aging occurs along general dimensions of variation in edge and node integrity suggests that these dimensions may be particularly relevant for cognitive aging. To test this hypothesis, we used the linear composite indices of connectome elements in LBC1936 (Fig. S2d), weighted by either PC loadings of connectome-wide edge or node integrity or age correlations in UKB, to test associations with latent processing speed, visuospatial ability, and memory factors (see Method). As would be expected on the basis of the sizable associations between age correlations and PC loadings, age-weighted and PC-weighted composites were very highly correlated (*r*_edge-based composites_ = −0.892; *r*_node-based composites_ = −0.999) and exhibited very similar patterns of associations with the cognitive outcomes. This indicates that brain age and overall integrity are virtually indistinguishable.

#### Edge-based composites

Composite indices of connectome-wide edge integrity were significantly associated with processing speed (*r*_age-weighting_ = −0.190; 95% CI = [-0.282, −0.099]; *r*_PC-weighting_ = 0.178; 95% CI = [0.085, 0.270]), but not with visuospatial ability (*r*_age-weighting_ = − 0.091; 95% CI = [-0.188, 0.006]; *r*_PC-weighting_ = 0.066; 95% CI = [-0.032, 0.163]) or memory (*r*_age-weighting_ = −0.083; 95% CI = [-0.186, 0.020]; *r*_PC-weighting_ = 0.053; 95% CI = [-0.051, 0.157]). For both age-weighting and PC-weighting, a 1000-fold permutation test (Fig. S9; Table S7; Supplementary Method & Results) in which the weights were randomly shuffled across edges indicated that observed edge-based composites were more predictive of both processing speed and visuospatial ability than over 99% of the permuted data (empirical *p*’s < 0.01) and more predictive of memory than over 95% of the permuted data (empirical *p*’s < 0.05).

Composite indices created for the individual NOIs varied in their magnitudes of prediction of processing speed (*r*_age-weighting_ range = −0.192 to −0.037; *r*_PC-weighting_ range = −0.099 to 0.186), but displayed null associations with visuospatial ability (*r*_age-weighting_ range = −0.153 to 0.009; *r*_PC-weighting_ range = 0.069 to 0.100) and memory (*r*_age-weighting_ range = −0.102 to 0.002; *r*_PC-weighting_ range = −0.034 to 0.100; top left panels of Figs. 5 & S10). Differences in the magnitudes of association across NOIs may stem from differences in their sizes, with large networks aggregating more information. To examine prediction relative to network size, we divided the magnitude of the correlation by the total number of elements on which the composite indices were each based (*r*_age-weighting_adjusted_ range = −0.0063 to −0.0001 with processing speed; *r*_PC-weighting_adjusted_ range = −0.0022 to 0.0066 with processing speed; top right panels of Figs. 5 and S10). Edge-based composite indices of Central Executive network integrity showed the largest size-adjusted magnitudes of association with processing speed. As edge strength was generally unrelated to visuospatial ability and memory, we do not interpret the size-adjusted associations with either domain of cognitive function.

**Figure 5.**
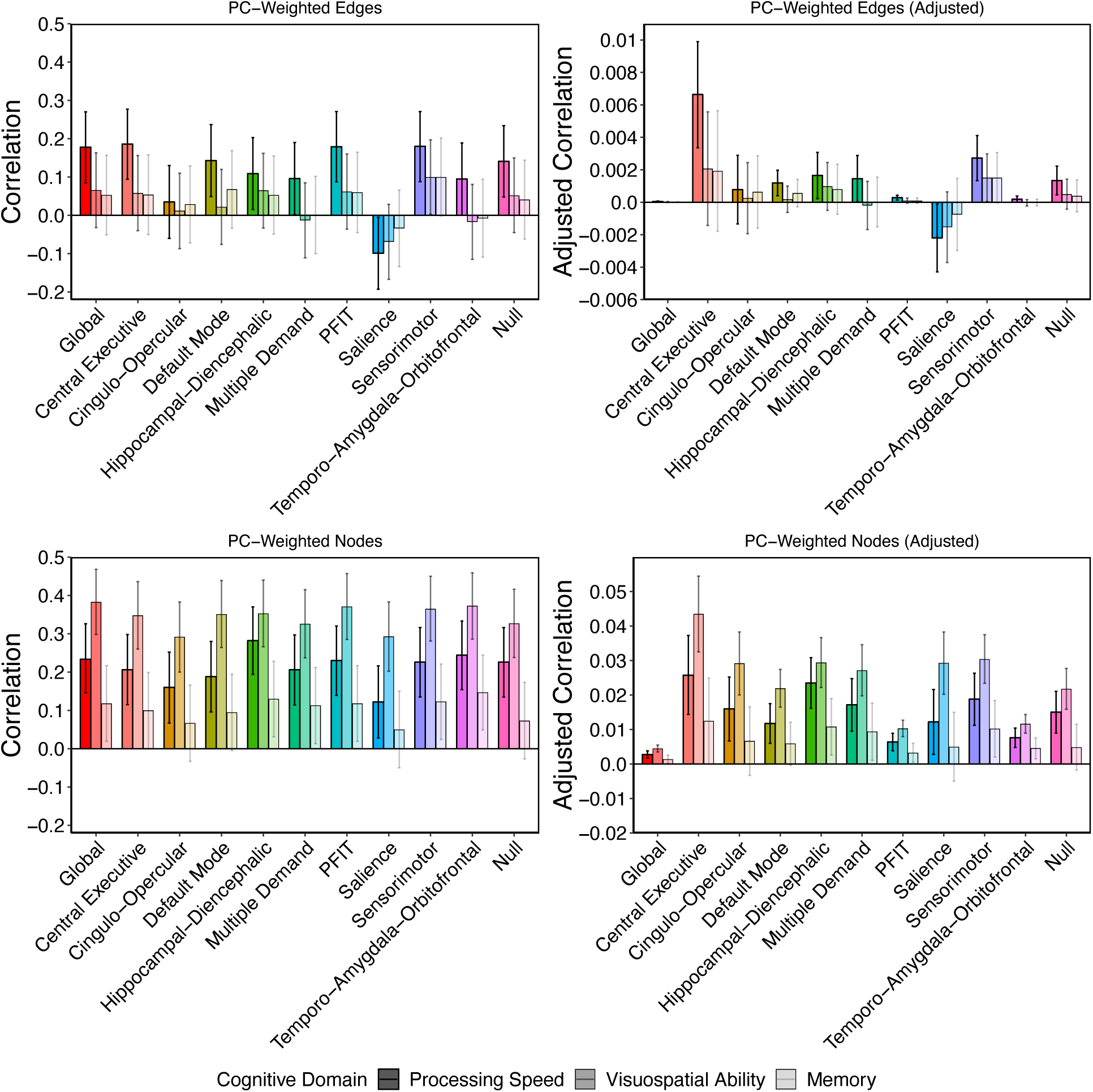
**Prediction of cognitive function from UKB-weighted indices of connectome integrity.** Raw and adjusted associations between weighted-composite scores reflecting variation in overall connectome integrity and cognitive function in LBC1936. Adjusted estimates were created by dividing the raw estimates by the number of edges or nodes in the network. Note that raw associations for edges and nodes are presented on the same *y*-axis scale, whereas the scale for the adjusted associations differs for edges and nodes. Scores were created across the whole brain and all NOIs by summing the LBC1936 data weighted by each element’s loading on the first principal component of its respective subnetwork discovered in UK Biobank. Plots are broken down by element type (*i.e.*, edges or nodes) and reflect correlations between respective weighted composites from each NOI and the cognitive domains of processing speed, visuospatial ability, and memory. Error bars represent 95% confidence intervals.

#### Node-based composites

Composite indices of connectome-wide node integrity were significantly associated with all cognitive domains (processing speed: *r*_age-weighting_ = −0.244; 95% CI = [-0.344, −0.154]; *r*_PC-weighting_ = 0.235; 95% CI = [0.146, 0.326]; visuospatial ability: *r*_age-weighting_ = −0.386; 95% CI = [-0.471, −0.301]; *r*_PC-weighting_ = 0.383; 95% CI = [0.298, 0.468]; memory: *r*_age-weighting_ = −0.124; 95% CI = [-0.223, −0.025]; *r*_PC-weighting_ = 0.118; 95% CI = [0.019, 0.217]). For both age-weighting and PC-weighting, a 1000-fold permutation test (Fig. S9; Table S7; Supplementary Method & Results) in which the weights were randomly shuffled across nodes indicated that observed node-based composites were not substantially more predictive of processing speed, visuospatial ability, or memory than the permuted data (empirical *p*’s > 0.08). This is consistent with the high intercorrelations among the node volumes, and the observations that the distributions of associations for nearly all permuted node runs were very narrow, indicating that node volumes may be largely exchangeable with respect to cognitive ability-relevant information.

Network-specific composite indices varied in their magnitudes of prediction across NOIs, with prediction of visuospatial ability generally exceeding that of processing speed or memory (processing speed: *r*_age-weighting_ range = −0.289 to −0.125; *r*_PC-weighting_ range = 0.122 to 0.282; visuospatial ability: *r*_age-weighting_ range = −0.376 to −0.281; *r*_PC-weighting_ range = 0.292 to 0.373; memory: *r*_age-weighting_ range = −0.151 to −0.062; *r*_PC-weighting_ range = 0.050 to 0.147; bottom left panels of Figs. 5 & S10). After adjusting for the number of elements, the age-weighted nodes in the Central Executive network displayed the largest magnitude of associations all domains of cognitive function (processing speed: *r*_age-weighting_adjusted_ = −0.026, 95% CI = [-0.038, −0.015]; *r*_PC-weighting_adjusted_ = 0.026, 95% CI = [0.014, 0.037]; visuospatial ability: *r*_age-weighting_adjusted_ = −0.044, 95% CI = [-0.055, −0.033]; *r*_PC-weighting_adjusted_ = 0.044, 95% CI = [0.033, 0.055]; memory: *r*_age-weighting_adjusted_ = −0.013, 95% CI = [-0.025, −0.0003]; *r*_PC-weighting_adjusted_ = 0.013, 95% CI = [0.000, 0.025]; bottom right panels of Figs. 5 & S10).

#### General dimensions of edge and node integrity predict late-life cognitive function incremental of TBV

We fitted multiple regression models in LBC1936 to test whether the associations between general dimensions of connectome integrity and cognitive function were unique of TBV, which is perhaps the most robust and well-validated structural MRI predictor of cognitive function^13, 17^. Results are presented in the top portions of each panel of Table 1. TBV displayed very strong associations with node-based composite scores (*r*_age-weighting_ = −0.866; *r*_PC-weighting_ = 0.876; all *p*’s < 0.0005), but weak associations with edge-based composites (*r*_age-weighting_ = −0.014, *p* = 0.750; *r*_PC-weighting_ = 0.017, *p* = 0.707).

**Table 1.** Associations between weighted connectome (edge and node) composites, total brain volume, and age 11 IQ.

| **Table 1a: Processing Speed** |  |  |  |  |  |  |  |
| --- | --- | --- | --- | --- | --- | --- | --- |
| *Composite* | *Model* | *Predictor 1* | *Predictor 2* | $\beta 1$ (p-value) | $\beta 2$ (p-value) | $R^2$ | *Multiple R* |
| **Age-based** | 1a | - | TBV | - | 0.165 (0.001) | 0.027 | 0.165 |
|  | 1b | Edges | TBV | -0.189 (< 0.0005) | 0.163 (0.001) | 0.063 | 0.251 |
|  | 1c | Nodes | TBV | -0.396 (< 0.0005) | -0.176 (0.062) | 0.067 | 0.259 |
|  | 2a | Edges only | - | -0.190 (< 0.0005) | - | 0.036 | 0.190 |
|  | 2b | - | Nodes only | - | -0.244 (< 0.0005) | 0.059 | 0.244 |
|  | 2c | Edges | Nodes | -0.162 (0.001) | -0.223 (< 0.0005) | 0.085 | 0.292 |
|  | 3a | - | Age 11 IQ | - | 0.511 (< 0.0005) | 0.261 | 0.511 |
|  | 3b | Edges | Age 11 IQ | -0.145 (0.001) | 0.498 (< 0.0005) | 0.281 | 0.530 |
|  | 3c | Nodes | Age 11 IQ | -0.161 (< 0.0005) | 0.484 (< 0.0005) | 0.285 | 0.534 |
| **PC-based** | 4a | - | TBV | - | 0.165 (0.001) | 0.027 | 0.165 |
|  | 4b | Edges | TBV | 0.175 (< 0.0005) | 0.163 (0.001) | 0.058 | 0.241 |
|  | 4c | Nodes | TBV | 0.379 (< 0.0005) | -0.165 (0.092) | 0.062 | 0.249 |
|  | 5a | Edges only | - | 0.178 (< 0.0005) | - | 0.032 | 0.178 |
|  | 5b | - | Nodes only | - | 0.234 (< 0.0005) | 0.055 | 0.234 |
|  | 5c | Edges | Nodes | 0.154 (0.001) | 0.219 (< 0.0005) | 0.079 | 0.281 |
|  | 6a | - | Age 11 IQ | - | 0.511 (< 0.0005) | 0.261 | 0.511 |
|  | 6b | Edges | Age 11 IQ | 0.134 (0.002) | 0.500 (< 0.0005) | 0.277 | 0.526 |
|  | 6c | Nodes | Age 11 IQ | 0.154 (< 0.0005) | 0.486 (< 0.0005) | 0.283 | 0.532 |

| **Table 1b: Visuospatial Ability** |  |  |  |  |  |  |  |
| --- | --- | --- | --- | --- | --- | --- | --- |
| *Composite* | *Model* | *Predictor 1* | *Predictor 2* | $\beta 1$ (p-value) | $\beta 2$ (p-value) | $R^2$ | *Multiple R* |
| **Age-based** | 1a | - | TBV | - | 0.333 (< 0.0005) | 0.111 | 0.333 |
|  | 1b | Edges | TBV | -0.084 (0.077) | 0.330 (< 0.0005) | 0.117 | 0.342 |
|  | 1c | Nodes | TBV | -0.395 (< 0.0005) | -0.011 (0.909) | 0.149 | 0.386 |
|  | 2a | Edges only | - | -0.091 (0.067) | - | 0.008 | 0.091 |
|  | 2b | - | Nodes only | - | -0.386 (< 0.0005) | 0.149 | 0.386 |
|  | 2c | Edges | Nodes | -0.039 (0.415) | -0.380 (< 0.0005) | 0.150 | 0.387 |
|  | 3a | - | Age 11 IQ | - | 0.553 (< 0.0005) | 0.306 | 0.553 |
|  | 3b | Edges | Age 11 IQ | -0.040 (0.372) | 0.549 (< 0.0005) | 0.307 | 0.554 |
|  | 3c | Nodes | Age 11 IQ | -0.308 (< 0.0005) | 0.504 (< 0.0005) | 0.397 | 0.630 |
| **PC-based** | 4a | - | TBV | - | 0.333 (< 0.0005) | 0.111 | 0.333 |
|  | 4b | Edges | TBV | 0.059 (0.219) | 0.331 (< 0.0005) | 0.114 | 0.338 |
|  | 4c | Nodes | TBV | 0.397 (< 0.0005) | -0.016 (0.869) | 0.146 | 0.382 |
|  | 5a | Edges only | - | 0.066 (0.187) | - | 0.004 | 0.066 |
|  | 5b | - | Nodes only | - | 0.383 (< 0.0005) | 0.147 | 0.383 |
|  | 5c | Edges | Nodes | 0.023 (0.628) | 0.380 (< 0.0005) | 0.147 | 0.383 |
|  | 6a | - | Age 11 IQ | - | 0.553 (< 0.0005) | 0.306 | 0.553 |
|  | 6b | Edges | Age 11 IQ | 0.019 (0.676) | 0.552 (< 0.0005) | 0.306 | 0.553 |
|  | 6c | Nodes | Age 11 IQ | 0.306 (< 0.0005) | 0.506 (< 0.0005) | 0.397 | 0.630 |

| *Table 1c: Memory* |  |  |  |  |  |  |  |
| --- | --- | --- | --- | --- | --- | --- | --- |
| *Composite* | *Model* | *Predictor 1* | *Predictor 2* | $\beta 1$ (*p*-value) | $\beta 2$ (*p*-value) | $R^2$ | *Multiple R* |
| **Age-based** | 1a | - | TBV | - | 0.012 (0.815) | 0.0001 | 0.012 |
|  | 1b | Edges | TBV | -0.083 (0.115) | 0.008 (0.877) | 0.007 | 0.084 |
|  | 1c | Nodes | TBV | -0.437 (< 0.0005) | -0.361 (< 0.0005) | 0.048 | 0.219 |
|  | 2a | Edges only | - | -0.083 (0.113) | - | 0.007 | 0.083 |
|  | 2b | - | Nodes only | - | -0.124 (0.014) | 0.015 | 0.124 |
|  | 2c | Edges | Nodes | -0.066 (0.212) | -0.166 (0.024) | 0.020 | 0.141 |
|  | 3a | - | Age 11 IQ | - | 0.613 (< 0.0005) | 0.376 | 0.613 |
|  | 3b | Edges | Age 11 IQ | -0.027 (0.551) | 0.615 (< 0.0005) | 0.381 | 0.617 |
|  | 3c | Nodes | Age 11 IQ | -0.033 (0.466) | 0.611 (< 0.0005) | 0.381 | 0.617 |
| **PC-based** | 4a | - | TBV | - | 0.012 (0.815) | 0.0001 | 0.012 |
|  | 4b | Edges | TBV | 0.053 (0.323) | 0.009 (0.866) | 0.003 | 0.055 |
|  | 4c | Nodes | TBV | 0.437 (< 0.0005) | -0.365 (< 0.0005) | 0.045 | 0.212 |
|  | 5a | Edges only | - | 0.053 (0.318) | - | 0.003 | 0.053 |
|  | 5b | - | Nodes only | - | 0.118 (0.020) | 0.014 | 0.118 |
|  | 5c | Edges | Nodes | 0.038 (0.480) | 0.114 (0.026) | 0.015 | 0.122 |
|  | 6a | - | Age 11 IQ | - | 0.613 (< 0.0005) | 0.376 | 0.613 |
|  | 6b | Edges | Age 11 IQ | -0.002 (0.969) | 0.616 (< 0.0005) | 0.379 | 0.616 |
|  | 6c | Nodes | Age 11 IQ | 0.028 (0.526) | 0.612 (< 0.0005) | 0.380 | 0.616 |

Note. TBV = Total brain volume

##### Processing speed

TBV was significantly associated with processing speed (*β* = 0.165, *p* = 0.001). However, edge- and node-based composites of connectome integrity predicted processing speed incremental of TBV (*edges*: *β*_age-weighting_ = −0.189; *β*_PC-weighting_ = 0.175; *nodes*: *β*_age-weighting_ = −0.396; *β*_PC-weighting_ = 0.379; all *p*’s < 0.0005).

##### Visuospatial ability

TBV was significantly associated with visuospatial ability (*β* = 0.333, *p* < 0.0005). However, node-based composites of connectome integrity predicted visuospatial ability incremental of TBV (*β*_age-weighting_ = −0.395; *β*_PC-weighting_ = 0.397; all *p*’s < 0.0005).

##### Memory

TBV was not significantly associated with memory (*β* = 0.012, *p* = 0.815). Node-based composites of connectome integrity predicted memory incremental of TBV (*β*_age-weighting_ = −0.437; *β*_PC-weighting_ = 0.437; all *p*’s < 0.0005)

#### General dimensions of edge- and node-integrity predict late-life cognitive function incremental of one another

We fitted multiple regression models to test whether the associations between edge- and node-based indices of connectome integrity and cognitive function were unique of one another. Results are presented in the middle portions of each panel of Table 1. All associations that were present in the univariate context remained in this multiple regression context. For processing speed, the multiple R^2^s from the models that included both edge- and node-based indices were over 40% larger than the R^2^s from models including only node-based indices, and over 100% larger than the R^2^s from models including only edge-based indices. For visuospatial ability and memory, multiple R^2^s from the models that included both edge- and node-based indices were only marginally larger than the R^2^s from models including node-based indices alone.

#### General dimensions of edge- and node-integrity predict late-life cognitive function incremental of childhood intelligence

The LBC1936 study has available a high-quality index of IQ at age 11 years, the Moray House Test No. 12. Age-11 IQ was associated with node-based indices of connectome integrity at age 73 (*β*_age-weighting_ = −0.159; *β*_PC-weighting_ = 0.155; all *p*’s <.0005) but was not significantly associated with edge-based indices of connectome integrity at age 73 (*β*_age-weighting_ = −0.080, *p* = 0.074; *β*_PC-weighting_ = 0.076, *p* = 0.090). These results are consistent with previous findings in the same sample that demonstrated comparable associations between age-11 IQ and other age-73 structural MRI indices (brain cortical thickness)^39^, and collectively suggest that general dimensions of node integrity may at least partially reflect lifelong brain health.

To probe whether associations between composite indices of age-73 connectome integrity and age-73 cognitive function were plausibly reflective of aging-specific processes, we examined whether the associations persisted after controlling for age-11 IQ. Results are presented in the bottom portions of each panel of Table 1. Age-73 connectome-integrity indices maintained their associations with age-73 processing speed and visuospatial ability even after controlling for age-11 IQ. The modest node-based associations with memory did not persist after controlling for age-11 IQ, suggesting that the association between age-73 node-based connectome integrity and age-73 memory function may be a vestige of associations between early-life differences in cognitive ability and node-based connectome integrity.

#### Regularized LASSO regression models

We were interested in whether a least absolute shrinkage and selection operator (LASSO) approach for indexing *connectome age* could improve prediction of late-life cognitive function beyond the simple composite indices reported above (see Supplementary Method for detail). LASSO models include penalty functions that shrink regression coefficients toward zero in order to introduce sparsity into the predictor set and guard against overfitting.

In UKB, a LASSO model based on all edges predicted 50.7% of the variance in age in the UKB holdout sample, whereas a model based on all nodes predicted only 33.4% of the variation in age (Fig. S11; Supplementary Results). The model based on edges alone had good prediction accuracy in a holdout subsample (RMSE = 5.32 years). The model based on nodes alone showed had slightly worse prediction accuracy than the model based on edges alone (RMSE = 6.1 years). A LASSO model based on both edges and nodes, and a LASSO model that incorporated Edge × Node interactions did not appreciably improve prediction of age relative to the model based on edges alone (Fig. S12; Supplementary Results).

Similar patterns of results were obtained for LASSO analyses based on NOIs, albeit at lower overall levels of age prediction (*R*^2^ = 0.166 to 0.507 using edges; 0.087 to 0.334 using nodes; Figs. S11 & S12), as would be expected when less information is made available for predictive modelling. Likewise, removing potentially spurious edges with consistency-based thresholding prior to conducting the LASSO analyses slightly depreciated predictions relative to unthresholded data (mean ratio of unthresholded *R^2^* to thresholded *R^2^* across NOIs = 1.477; Supplementary Results).

We predicted cognitive function in LBC1936 from connectome elements, using LASSO models trained in UKB (see Supplementary Method). We confine our prediction to processing speed and visuospatial ability, as memory was not associated with connectome integrity above and beyond age-11 IQ. Edge-based LASSO predictors were generally not significantly related to cognitive function (Fig. S13). Node-based LASSO predictors were significantly associated with both processing speed (*r*_whole-brain_ = −0.199; 95% CI = [-0.289, −0.109]) and visuospatial ability (*r*_whole-brain_ = −0.177; 95% CI = [-0.271, −0.083]; Fig. S13). Effect sizes were not appreciably larger than those obtained using the simple composite indices (outside of a LASSO framework) reported earlier (Fig. S14). Thus, the sparsity introduced by regularized methods was not advantageous – and in the case of edge-based predictors, was disadvantageous – for predicting variation in late-life cognitive abilities from indices of connectome aging. Rather, the simple regression-weighted MRI composite scores reported earlier are able to produce impressively large associations with cognitive function that rival, if not exceed, those obtained for more complex algorithmic learning methods^40^.

## Discussion

Elucidating the neural bases of cognitive aging will be fundamental to detecting, and ultimately mitigating, preventing, or ameliorating aging-related cognitive impairments. Rather than focus on TBV, or a narrow selection of very specific regions of interest, we undertook a network approach that examined variation in elements of the whole-brain connectome and several of its NOIs in relation to variation in adult chronological age and adult cognitive abilities. Using age-heterogenous data from UKB, we found that the aging of elements within the connectome occurs along the same general dimensions of global brain health that underlie age-partialled correlations amongst connectome element integrities. We used indices of these general dimensions of edge and node integrity in LBC1936 to predict processing speed, visuospatial ability, and memory, three aging-sensitive domains of cognitive function in older adulthood^8,9,10^. Indices of connectome-wide node integrity were related to all domains of cognitive function, whereas indices of connectome-wide edge integrity were specifically related to processing speed. Network-specific analyses indicated a disproportionally large role of the Central Executive network in these patterns relative to its small size. Associations with processing speed and visuospatial ability were incremental of TBV, and persisted after controlling for age-11 IQ, suggesting that they capture aging-specific processes, whereas associations with memory appear to be a vestige of early-life differences in cognitive function.

Our analysis within UKB revealed a potentially important connection between individual differences in neurostructural integrity and aging-related decrements. We found that connectome elements that had stronger loadings on their corresponding PCs (i.e., elements that were better indicators of overall levels of neurostructural integrity) also tended to have stronger correlations with age. Although loadings on the PCs were robustly related to the centrality of each element within the physical topology of the connectome, this association was not sufficient to explain the strong correspondence between PC loadings and age correlations of the individual connectome elements. This pattern parallels results from cognitive aging research, where similar methodological approaches have found that tests with stronger loadings on a general factor of cognitive ability, indicated by many different measures, also tend to be more closely correlated with age^41, 42^, suggesting a strong shared genetic basis for cognitive aging across different ability domains^43^. The current results extend this phenomenon to the context of the brain and suggest that researchers addressing the causes of individual differences in aging-related cognitive and neurostructural decline would benefit from focusing their efforts on understanding broad, general mechanisms of aging, in addition to more specific or granular mechanisms.

That brain aging occurs along general dimensions of individual differences in connectome integrity suggests that the neurobiological substrates of cognitive aging may be very broad, but also raises considerable interpretation challenges to work on apparent brain age. Apparent brain age may simply be a marker of overall connectome health or integrity, rather than a marker specific to apparent aging. Using a high-quality measure of age-11 IQ, we found that late-life connectome integrity is partly accounted for by pre-existing differences in cognitive ability prior to the initiation of aging. By controlling for age-11 IQ, we confirmed that the detected associations between age-73 connectome integrity and age-73 processing speed and visuospatial ability are also likely to be partly reflective of the aging process proper. Other work that does not have high-quality controls for prior intelligence and/or brain structure will need to exercise caution when interpreting associations between indices of brain age and external outcomes. Not only were age correlations strongly related to PC loadings, but age-correlation weighted indices of connectome age were nearly entirely collinear with PC-loading weighted indices of connectome integrity (*r*_edge-based composites_ = −0.892; *r*_node-based composites_ = −0.999). Thus, any given association with apparent brain age might just as appropriately be conceptualized as an association with overall brain integrity.

It is notable that elements within the Salience network were found to be relatively spared in late-life, implying that its trajectory across middle to older adulthood is flatter than other NOIs. Previous functional research has suggested that the Salience network acts as a control on other networks, such as the Central Executive and Default Mode networks, and that disruption of this control, or loss of functional connections between networks, is among the causes of cognitive decline^44,45,46^. Though we did not examine between-network connections here, our analyses did highlight the Central Executive network as being of particular interest for cognitive aging, despite it containing the fewest number of elements of any network. To correct for the additive effect of the size of each network, we adjusted their predictions of age and cognitive functions for their respective number of elements. Consistently across methodological approaches, element types, and outcome variables, the Central Executive network stood out after this adjustment: its small number of elements had disproportionately stronger associations with both processing speed and visuospatial ability relative to its size. Thus, the integrity of this network is likely to be of particular relevance to cognitive aging.

Although this study examined an important, theoretically-informed set of brain networks in large-scale samples with high-quality brain-imaging and cognitive testing, and used a cross-cohort (wide-to-narrow age range) design to limit problems of overfitting, it nevertheless had some key limitations. First, though the LBC1936 (testing) and UKB (training) samples were non-overlapping, they still had many qualities in common: they were based in the United Kingdom, had the same broad ethnic and cultural background, and – perhaps most importantly – they were self-selecting samples that were healthier, better-educated, and more cognitively able than average^47, 48^. It would be of interest to examine whether the brain-network predictors derived here are still effective in samples that are further removed from this context, or are more representative of the broader population. For this reason, we have made the weightings for each of our predictors publicly available for the use of other researchers in their own appropriate datasets (see Table S9). Second, the study focused on predicting cross-sectional differences in cognitive function in old age from connectome indices alone. Future work would benefit from investigating neural predictors of the longitudinal slopes of cognitive change in late-life, assessing whether the brain networks that explain individual differences in cognitive level are the same as those that explain individual differences in cognitive decline. Relatedly, the predictors of cognitive function were trained on cross-sectional differences in brain structure. Research integrating measurement of aging-related brain changes with previously identified determinants of cognitive decline^49^, including medical comorbidities (e.g., small vessel disease indicators, inflammation, vascular disease), lifestyle indicators (e.g., diet, smoking, physical function), and genetic risk, may help to critically advance prediction of cognitive aging. Third, our analyses were based on unthresholded connectivity matrices. Though we found that edge-wise age trends and PC loadings were largely unchanged by thresholding, it is possible that edges that occur in very few subjects and involve very few streamlines contain greater measurement error^50,51^. Fifth, the LBC1936 and UKB MRI scanners differed in acquisition strength (1.5T and 3T, respectively). It is potentially nontrivial to compare brain indices across scanners of different magnetic strength, and future research would benefit from assessing the extent to which these differences bias results in cross-cohort MRI studies. Finally, the neurostructural perspective is inherently limited in that it does not include functional information; since previous studies have found functional connections between several of the networks studied here^44^, our investigation, which treated networks separately, may have missed these links, which might explain additional cognitive variance over and above the properties of the individual networks. Integrating the structural and functional perspectives is a critical future task for network-focused cognitive neuroscience.

This study represents the most comprehensive investigation to date of the out-of-sample predictive validity of several theoretically-informed brain structural networks for late-life cognitive function. We found evidence that aging in the brain as a whole, and within specific networks, tends to occur on broad, general dimensions, with brain features that are more representative of their network in general being more related to age. Age-related elements of each network often made substantial out-of-sample predictions of cognitive abilities, with the Central Executive showing disproportionately large relation with processing speed, visuospatial ability, and memory relative to its size. Given the wealth of neuroimaging data now available, the cross-cohort-comparison approach will be a viable and fruitful way of producing predictors of cognitive abilities that are robust to context, and thus of potential use in predicting and understanding differences in cognitive aging.

## Acknowledgments

This work was supported by National Institutes of Health (NIH) grant R01AG054628. The Population Research Center at the University of Texas is supported by NIH grant P2CHD042849. The Lothian Birth Cohorts group is funded by Age UK (Disconnected Mind grant), the Medical Research Council (grant MR/R024065/1), and the University of Edinburgh’s School of Philosophy, Psychology and Language Sciences. This research was conducted using the UK Biobank Resource (Application Nos. 10279). We thank the UK Biobank participants and UK Biobank team for their work in collecting, processing, and disseminating these data for analysis.

## Author contributions

SJR, EMT-D, & JWM conceived of the study design. CRB & SRC processed the UKB MRI data. SRC, MAH, & MVH processed the LBC1936 MRI data. JWM and SJR conducted the analyses under supervision from EMT-D. JWM, SJR & EMT-D wrote the paper. All authors contributed edits and feedback on the paper.

## Data Availability

UKB-derived age- and PC-weights for connectome elements are available as Table S9.

## Methods

### Participants

#### UK Biobank

A large-scale population epidemiology study, UK Biobank (UKB) involved the recruitment of approximately 500,000 individuals across Great Britain for medical, psychosocial, and biological data collection^52^. A subset of around 100,000 UKB participants were invited to complete brain MRI scanning (scanner details are provided in the next section); as of this writing, data collection is still in progress, but portions of the data have been made available. The initial release of diffusion MRI (dMRI) data included 5,455 individuals. Data for a subset of these individuals (*n* = 567) was acquired at an earlier scanning phase, rendering their dMRI data incompatible with subsequent data acquisition (see section 2.10 of the Brain Imaging Documentation (http://biobank.ctsu.ox.ac.uk/crystal/refer.cgi?id=1977) for details). A further subset (*n* = 1,314) was removed during dMRI quality-control procedures prior to data release. In the present study, 3,155 participants (1,623 female) who had MRI data were included, with an average age of 61.6 years (SD = 7.5, range = 44.64 – 77.12). Of the 3,155 participants with structural MRI data, 3,124 had usable volume data and 3,087 reported their age at the time of scanning. All participants were free of potentially confounding dementias and neurological syndromes (e.g., multiple sclerosis, stroke). Despite previous research that has demonstrated neuroanatomical sex differences in men and women^20, 53^, we found largely similar patterns of connectome aging across men and women (*r*_edge-age correlations_ = 0.852; *r*_node-age correlations_ = 0.943, all *p*’s < 0.0005). We therefore did not further correct for biological sex. All the data from the present study come from the UK Biobank recruitment center in Manchester, UK. UKB received ethical approval from the Research Ethics Committee (reference 11/NW/0382). All participants provided informed consent to participate.

#### Lothian Birth Cohort 1936

In 1947, almost all children attending schools in Scotland and born in 1936 completed an intelligence test as part of the Scottish Mental Survey 1947^54^. 1,091 of these individuals living mostly in the Edinburgh and Lothians area of Scotland were contacted and returned for re-testing at an average age of 69.5 years, becoming the Lothian Birth Cohort 1936 (LBC1936)^48, 55^, a longitudinal study of aging. As part of the second wave of testing, at age 72.8 years (SD = 0.70), 731 LBC1936 members underwent brain MRI scanning (see scanner details below), of whom 528 (246 female) had reliable brain and cognitive data for the cognitive prediction analysis. Participants were largely healthy: only seven scored in the mild range of dementia on the Mini-Mental State Exam, zero self-reported symptoms of dementia, and 65 met for neuroradiologically-identified stroke^56^. Only data from this second wave are included in the present study.

### Brain Image Acquisition and Processing

#### UK Biobank

MRI data for all participants was collected on the same 3T Siemens Skyra MRI scanner (see Miller et al.^57^& Alfaro-Almagro et al.^58^ for full details). T1-weighted volumes were acquired in the sagittal plane using a 3D MP-RAGE sequence. The T1-weighted volumes were preprocessed and analyzed using FSL tools (http://www.fmrib.ox.ac.uk/fsl) by the UKB brain imaging team. A full overview of the preprocessing and analysis pipeline is available at http://biobank.ctsu.ox.ac.uk/crystal/docs/brain_mri.pdf. FoV-reduced T1-weighted volumes from the first release of UKB MRI data were used to reconstruct and segment the cortical mantle using default parameters in FreeSurfer v5.3^59^ (http://surfer.nmr.mgh.harvard.edu/). Reconstruction and segmentation were based on the Desikan-Killiany atlas^36^. Automated anatomical segmentation of subcortical structures (accumbens area, amygdala, caudate, hippocampus, pallidum, putamen, thalamus, ventral diencephalon, and brainstem) was also conducted in FreeSurfer using default settings and atlas^60^. Each output underwent visual assessment and subjects were excluded if major errors in tissue identification or skull stripping were identified (which were not analyzed further).

#### Lothian Birth Cohort 1936

MRI data for all participants was collected at Wave 2 on the same GE Signa Horizon HDx 1.5T clinical scanner (General Electric, Milwaukee, WI) equipped with a self-shielding gradient set (33 mT/m maximum gradient strength) and manufacturer supplied eight-channel phased-array head coil (see Wardlaw et al.^56^ for full details). High-resolution T1-weighted volumes were acquired in the coronal plane using a 3D fast-spoiled gradient echo (FSPGR) and subsequently processed in FreeSurfer v5.1. As with the UK Biobank data, reconstruction and segmentation were based on the Desikan-Killiany atlas^36, 60^. Segmentation and parcellation errors were corrected manually after visual inspection of each image.

#### Tractography

Probabilistic tractography pipelines were identical across both datasets, though acquisition procedures differed slightly. For UKB, dMRI acquisitions are publicly available from the UKB website in the form of a Protocol (http://biobank.ctsu.ox.ac.uk/crystal/refer.cgi?id=2367), Brain Imaging Documentation (and in Miller et al.^57^)). The dMRI data were acquired using a spin-echo echo-planar imaging sequence (50 b = 1000 s/mm^2^, 50 b = 2000 s/mm^2^ and 10 b = 0 s/mm^2^) resulting in 100 distinct diffusion-encoding directions. The field of view was 104 × 104 mm with imaging matrix 52 × 52 and 72 slices with slice thickness of 2 mm resulting in 2 × 2 × 2 mm voxels. For LBC1936, dMRI data was acquired from both T2-weighted and sets of diffusion-weighted (b = 1000 s/mm^2^) axial single-shot spin-echo echo-planar (EP) volumes acquired with diffusion gradients applied in 64 noncolllinear directions^56^. Both datasets were corrected for head motion and eddy currents, and processed using BEDPOSTx, with within-voxel modeling of multi-fibre tract orientation structure. Probabilistic tractography with crossing fiber modeling was carried out using PROBTRACKx^61^. Streamlines were seeded from all white matter voxels using 100 Markov Chain Monte Carlo iterations with a fixed step size of 0.5 mm between successive points.

#### Connectome Construction

Our treatment of the structural brain data from both UKB and LBC1936 is based on an automated connectivity mapping pipeline^62, 63^, wherein T1-weighted volumes are decomposed into 85 distinct cortical and subcortical nodes based on the Desikan-Killiany atlas. Such segmentations are used to model the brain as a structural network (i.e., *connectome*) comprised of nodes, or variables in the network, and edges, or the connections between nodes. As such, we constructed connectomes for each participant in UKB and LBC1936, where nodes represented grey matter regional volumes and white matter edge weights were the mean fractional anisotropy averaged along the length of all streamlines identified between each pair of nodes (*k* = 3,570 possible edges). Fractional anisotropy is a diffusion tensor MRI-derived measure of white matter organization that describes the directional coherence of water molecule diffusion. Six edges were estimated as zero across all participants (i.e., probabilistic tractography found no route between the nodes involved). Network data were not thresholded. For graph-theory analyses, we used the *igraph*^64^ package in R to compute network parameters.

### Networks-of-Interest

In order to develop greater specificity of the neuroanatomical correlates of cognitive aging, we decomposed each participant’s whole-brain connectome into ten prespecified NOIs (see Fig. 1; Table S1). NOIs were largely representative of previous, intrinsically-defined networks in the human brain^24, 25^, several of which have been previously implicated in cognitive ability (Table S1). A null network was created from regions that were not included in any of the prespecified NOIs.

### Cognitive Testing

Members of the LBC1936 have completed a range of cognitive tests at every wave of testing. For the present study, we focused on tests from the domains of processing speed, visuospatial ability, and memory, which have been characterized within this cohort in our previous research^65^. Visuospatial ability was measured using tests of Matrix Reasoning and Block Design from the Wechsler Adult Intelligence Test, 3^rd^ UK Edition (WAIS-III^UK^)^66^, and the Spatial Span test (forwards and backwards) from the Wechsler Memory Scale, 3^rd^ UK Edition (WMS-III^UK^)^67^. Processing speed was measured using the Digit-Symbol Substitution and Symbol Search tests from the WAIS-III^UK^, a test of 4-choice reaction time administered on a dedicated button-box instrument^68^, and a psychophysical test of inspection time administered on a computer monitor with a fast refresh rate^69^. Memory was measured using the Logical Memory and Verbal Paired Associates subtests of the WMS-III^UK^ and the Digit Span Backward subtest of the WAIS-III^UK^. All cognitive domains were modeled as latent variables. Fit indices, factor model parameter estimates, and descriptive statistics for the cognitive tests are reported in Table S8.

### Statistical software

All analyses were run in R^70^. Graphics were created using the *ggplot2* package^71^. Anatomical network plots were created using BrainNet Viewer^72^. Factor modeling and structural regression models were estimated using the *lavaan* package^73^. LASSO model fitting and associated cross-validation was conducted within the *cv.glmnet* package^74^, with wrapper functions from the *caret* package^75^ in R (see Supplementary Method for details).

### Code availability

Sample code to run analyses presented here, including discovery of age correlations and principal component loadings in UKB, creation of weighted composite scores in LBC1936, and latent variable associations with cognitive function in LBC1936 will be made available upon publication.

