## Supplementary Materials for "Enhanced prediction of cognitive function using aging-sensitive networks within the human structural connectome"

### 1. Supplementary Method

**a. Sensitivity analyses.** To probe the uniqueness of signatures of whole-brain and network-specific connectome integrity in predicting cognitive function, we ran a series of sensitivity analyses.

*i. Permuted composites.* We were interested in assessing whether the observed association between indices of connectome integrity and cognitive function were superior to those that would have been obtained had the weights used to create the indices been shuffled. We compared the observed associations between composite scores and all domains of cognitive function to associations with composite scores weighted by randomly-shuffled age correlations or PC loadings in the whole-brain. We shuffled weights for both edges and nodes in UKB (*k* = 1000 for each) by sampling with replacement from the observed age correlations or PC loadings (see Fig. 4). We then weighted and summed the LBC1936 connectomes by the shuffled weights, resulting in four random composite scores (edges and nodes weighted by either shuffled age correlations or shuffled PC loadings) for each participant. We calculated the association between processing speed, visuospatial ability, and memory and each of the 1000 permuted composites to arrive at an empirical null distribution. Observed associations were determined to be significant if they were smaller than the bottom 2.5%-ile or larger than the top 97.5%-ile of the empirical null distribution.

*ii. Thresholding analyses.* We probed whether associations between indices of connectome integrity and cognitive function were likely to be inflated by potentially spurious connections arising from the use of unthresholded data. In UKB, we re-estimated each connectome using a consistency-based thresholding approach^1^, wherein connections are retained only if they have sufficiently low inter-subject variability in their weight and if their weight is plausibly strong for its length, which has been suggested as a method for removing connections that are most likely to be spurious. We correlated the age correlations and the PC loadings estimated in the unthresholded connectomes, either when the thresholded edges were set to 0, or when they were deleted entirely (2601 edges retained).

**b. Regularized LASSO Regression Models.** We first trained models to predict age in UK Biobank, and used the resulting predictive model to compute indices of connectome age in LBC1936, which we used to predict cognitive function.

*i. Age prediction in UK Biobank.* We fitted LASSO models predicting chronological age using connectome elements, as well as dummy variables representing the presence or absence of an element, in 80% of the UKB sample (~2,520 participants), with 10-fold cross-validation to select the optimal penalization parameter (λ) that provided the lowest prediction error. We then used the obtained coefficients to produce a score on *connectome age* in the 20% UKB holdout sample (~630 participants), which we used to predict chronological age. This process was repeated 100 times – reported coefficients were averaged across each of the 100 runs. For the whole-brain connectome, and for each network-of-interest (NOI), we ran analyses separately for edges and nodes, with all edges and nodes together, and with a novel topologically-constrained weighting scheme that takes into account the interactive effects of edges and nodes.

This procedure was run for each NOI, separately for the network’s edges, nodes, and then with multiple weighting schemes reflecting the joint contribution edges and nodes together (see below for description). For edges, we included both the FA-weighting of each edge, as well as a binary dummy variable indicating whether or not the edge was present in a given participant.

Each iteration split a different random shuffle of the full dataset into training and testing samples. The results reported below for the variance explained in age by each network are the average *R*^2^ values from this resampling procedure, and the standard error for each *R*^2^ is the standard deviation of the values across all 100 iterations. We also report the *R*^2^ adjusted by the size of the network by dividing by the number of elements in the included predictor set.

*ii. Cognitive function prediction in LBC1936.* For the final predictive analysis, we employed a similar pipeline to train for predicting cognitive function in LBC1936 from connectome elements, using LASSO models trained in UKB. We first fitted LASSO models predicting chronological age using connectome elements in UKB participants, with 10-fold cross-validation to select the optimal penalization parameter (λ). We then used the obtained coefficients to produce a score on *connectome age* in the narrow-aged LBC1936 sample, which we used to predict processing speed and visuospatial ability. Note that, because participants were virtually identical in their age in LBC1936, any differences in their *connectome age* could not be attributable to actual differences in chronological ages.

*iii. Alternative weighting schemes.* To investigate whether edges and nodes, when considered jointly, were more strongly predictive of connectome or cognitive aging, we ran the LASSO-regression prediction analyses using elements from three joint-element weighting schemes: node + edge, node*edge, node*edge + node.

**Node + edge**. First, we sought to examine the contribution of nodes and edges considered additively. Under this weighting scheme, we included the raw elements from the structural connectome: node volumes (N=85) and FA-weighted edges (N=3570). This weighting scheme thus allows us to prune our estimates of potentially redundant information provided *across* nodes and edges.

**Node*edge**. Next, we sought to examine the holistic interaction of nodes and edges as tripartite systems. That is, we were interested not only in how nodes and edges collectively contributed to connectome and cognitive aging, but how their interactions predicted these outcomes. The steps of this weighting scheme, with accompanying mathematical representations, are detailed below. To summarize, we began by creating a vector of the square root of the 85 node volumes for each participant (*n* = 3,058 with full regional volume and age data). We then multiplied this vector by its transpose to create an 85x85 matrix of pairwise node weights in each participant. The off-diagonal elements in this matrix therefore encapsulate information about node integrity for pairwise combinations of nodes. Finally, we element-wise multiplied the pairwise node-weighted matrices by the edge-weighted (i.e., fractional anisotropy) matrices to create an 85x85 node*edge-weighted matrix in each participant. Elements in the resulting matrix are thus reflective of the strength of the connection between two nodes, as well as the structural integrity of these nodes. Our focus was on multiplicative, rather than additive, interactions between edges and nodes as we were interested in the holistic integrity of each pair of nodes and the connections between them, rather than merely the sum of their constituent parts. For example, if Node A has a robust volume and is strongly connected to Node B, but Node B is highly atrophied, then a coefficient measuring the systemic integrity of this neural pathway should adequately reflect the depreciable contribution of Node B to the system. That is, small values within a tripartite system should nullify larger values as we are interested in the unified structure of these systems. Thus, these matrices were used to estimate a network wherein edges represent not just white matter connections between regions, but the multiplicative effect of (sub)cortical volume by white matter connection strength.

**Step 1**: Create a vector of the square root of each node volume (N = 85) for each participant.

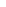

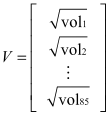

**Step 2**: Multiply the vector of node volumes by its transpose to create an 85x85 matrix, where cells represent the product of pairwise volumes.

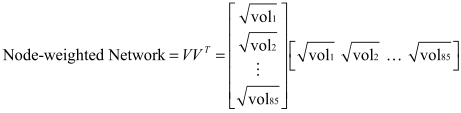

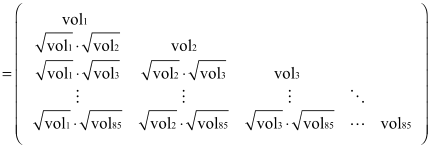

**Step 3**: Assemble the 85x85 edge-weighted matrix, where cells represent the microstructural connections between pairwise nodes.

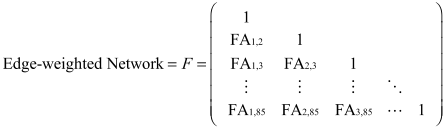

**Step 4**: Element-wise multiply the node-weighted matrix by the edge-weighted matrix to create an edge & node weighted matrix, where cells represent the multiplicative interaction between pairwise node volumes and the connection strength between those nodes.

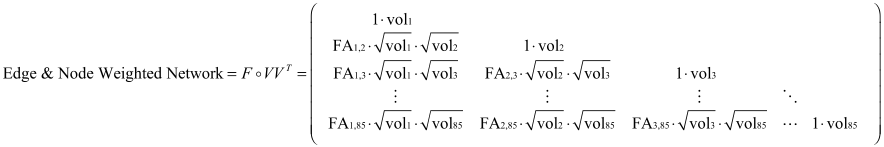

**Node*edge + node**. Lastly, we were interested in assessing whether the node*edge weighting scheme contributed more predictive power when nodes were included. Theoretically, if all useful information provided by nodes was captured in the node*edge weights, then the contribution of node volumes would have been redundant and dropped during the regularization process. Like the node + edge weighting scheme, this weighting scheme includes the 85 node volumes and the 3570 node*edge weights.

### 2. Supplementary Results

**a. Intercorrelations between network-specific PC-weighted composite scores.** We examined associations between composite indices of integrity within each NOI. Specifically, we computed intercorrelations amongst the network-specific PC-weighted composite scores for both edges and nodes to assess whether the general dimensions of network-specific integrity reflected an even broader dimension of whole-brain integrity (Fig. 2d; Fig. S4). Within edge-based composites, the average intercorrelation was 0.527 (interquartile range = 0.385 to 0.688; Table S5), with the first PC accounting for 59.7% of the variation in edge-based composite score covariance. Within node-based composites, the average intercorrelation was 0.812 (interquartile range = 0.746 to 0.872; Table S5), with the first PC accounting for 83.5% of variation in node-based composite score covariance. Edge-based composite scores and node-based composite scores were relatively uncorrelated within networks (see Table S6). Thus, after aggregating individual elements together according to prespecified NOIs, we observed strong correlations across the edge and node integrities of the different subnetworks. This is likely to be attributable both to the aggregation signal across elements within NOIs, that is itself correlated across NOIs, and to the overlap of elements within NOIs.

**b. Connectome aging occurs along general dimensions of variation in edge and node integrity: Analyses within NOIs.** Edges from several NOIs showed patterns consistent with the whole brain (Figs. S5 & S6). In the Central Executive, Cingulo-Opercular, Default Mode, Multiple Demand, PFIT, and Temporo-Amygdala-Orbitofrontal networks, edges with higher PC loadings tended to have stronger negative correlations with age (*r*s = -0.809 to -0.500). The Sensorimotor network was the only NOI to show a moderately strong positive association between PC loadings and age correlations (*r* = 0.312), suggesting that edges that are more central to the integrity of this network tend to be less susceptible to ageing-related degradation.

On average, within-NOI node results were similar to within-NOI edge results. Across several NOIs, there was a negative relationship observed between age correlations and PC loadings (*r*s = -0.937 to -0.352 in the Cingulo-Opercular, Default Mode, Hippocampal-Diencephalic, Multiple Demand, PFIT, Salience, and Temporo-Amygdala-Orbitofrontal networks). It is important to note that in certain networks (e.g., the Central Executive network), all elements had consistently strong PC loadings *and* age correlations (age *r*s < -0.189; PC loadings (λs > 0.69)), resulting in a weak association between the PC loadings and age correlations (*r* = -0.008).

**c. Associations between topological centrality, PC loadings, and age correlations in UKB.** To quantify the weighted connectedness of each node in the whole-brain network, we calculated its topological *strength* (i.e., sum of adjacent edge weights), which we then averaged across participants. To quantify the weighted connectedness of each edge in the whole-brain network, we calculated the average topological strength (i.e., average sum of adjacent edge weights) of the two nodes connected by that edge, which we then averaged across participants. We found that the topological strengths of both edges and nodes were strongly correlated with loadings on their respective PCs (*r* = 0.650 in edges; *r* = 0.558 in nodes; all *p*’s < 0.0005; see Fig. S7), suggesting that more topologically central elements within the connectome are more broadly representative of individual differences in the integrity of those elements. Importantly, this association is not due to a mathematical dependency between topological strength and principal component loadings, as topological strength is based on the absolute value of the edge weights whereas PC loadings are based on their covariation. The topological strength of an element was only modestly associated with its age correlation (*r* = -0.194 for edges, *p* < 0.0005; *r* = -0.235 for nodes, *p* = 0.031; see Fig. S8). Topological centrality is therefore insufficient to explain the observed associations between PC loadings and age correlations. In other words, connectome aging occurs along general dimensions of *variation* in edge and node integrity, but occurs only modestly, and proportionally, to the amount of topological connectedness of structural connectome elements.

**d. Results of regularized LASSO regression models.**

*i. Prediction of age in the UKB hold-out sample.* Figure S11 displays the raw and adjusted *R*^2^ values for each network for the LASSO regression prediction of age in the UK Biobank hold-out sample. Across the whole brain and all networks, edges alone explained greater variance in age than nodes alone (*R^2^* = 0.166 to 0.507 in edges; 0.087 to 0.334 in nodes). Edges from the PFIT and Temporo-Amygdala-Orbitofrontal networks accounted for the greatest variation in age (*R^2^* = 0.357, 95% CI = [0.312, 0.402]; *R^2^* = 0.277, 95% CI = [0.228, 0.326], respectively). For edges, only five dummy variables indicating the presence or absence of an edge were retained in 100% of the runs. There was only a modest boost when considering edge integrities (edges retained in 80+% of the runs = 3.1%). For nodes, each of the 85 node integrities was retained in at least 15% of the runs, with 44.7% of integrities appearing in 100% of the runs.

Adjusted estimates indicated that nodes explained substantially greater variance in age than edges (*R^2^* = 0.0039 to 0.0140 for nodes; *R^2^* = 0.0001 to 0.0060 for edges). Similar to the weighted-composite analyses, edges from the whole brain, PFIT and Temporo-Amygdala-Orbitofrontal networks explained substantially less variance in age than all other networks (*R^2^* = 0.0001 to 0.0006, 95% CIs = [0.0001, 0.0006]). Likewise, nodes from the whole brain and the PFIT network explained less variance than all but the Multiple Demand network (*R^2^* = 0.039 to 0.041, 95% CIs = [0.0033, 0.0053]). Overall, when adjusted for the number of elements in the network, the nodes provided more substantial age prediction.

Removing potentially spurious edges with consistency-based thresholding prior to conducting the LASSO analyses slightly depreciated predictions relative to unthresholded data (mean ratio of unthresholded *R^2^* to thresholded *R^2^* across NOIs = 1.477), though this was largely driven by sizeable differences in the Null (mean ratio of unthresholded *R^2^* to thresholded *R^2^* = 2.057) and Salience networks (mean ratio of unthresholded *R^2^* to thresholded *R^2^* = 3.311). To note, the correlation between mean edge weight and age within the Salience network remained positive, even after removing thresholded edges (*r* = 0.044, *p* = 0.012), indicating that the reported positive age trends in the Salience network are not simply an artifact of potentially spurious connections. Looking across all edges, however, prediction with thresholded data was nearly identical to prediction with unthresholded data, suggesting the utility of LASSO in zero-weighting uninformative edges (whole-brain *R^2^* = 0.505; ratio of unthresholded *R^2^* to thresholded *R^2^* in whole-brain = 1.004).

*ii. Prediction of cognitive function in LBC1936.* Figure S13 displays the raw and adjusted prediction results for processing speed and visuospatial ability. In contrast to the age-prediction results, nodes were substantially more predictive of both domains of cognitive function than were edges. For processing speed, node-based connectome age across the whole brain and almost all networks displayed relatively consistent associations (*r*s = -0.260 to -0.171, except in the Salience network, where *r* = -0.014). For visuospatial ability, nodes demonstrated somewhat more varied associations, with the Central Executive, Default Mode, Multiple Demand, PFIT, and Sensorimotor networks all displaying *r*’s stronger than -0.300. Such correlation magnitudes are comparable to the associations using the weighted composite scores (Fig. 5), demonstrating that prediction of cognitive function using grey-matter elements is retained in the context of a regularization approach that favors sparsity of the predictor set. Edge-based connectome age, however, displayed relatively weak associations with both processing speed and visuospatial ability (*r*’s = -0.119 – 0.125; -0.090 to 0.037 for the two abilities respectively). These associations were somewhat lower than the weighted composite scores, indicating that the statistical-learning algorithm did not retain the same predictive ability for cognitive function.

Consistent with the composite analyses, node-based brain age from the Central Executive network retained the strongest adjusted correlations with both processing speed and visuospatial ability of any network (*r*_adj_ for processing speed = -0.0319, 95% CI = [-0.0431, -0.0206]); *r*_adj_ for visuospatial ability = -0.0429, 95% CI = [-0.0539, -0.0319]). Across both processing speed and visuospatial ability, the Sensorimotor, Multiple Demand, and Cingulo-Opercular networks demonstrated the strongest adjusted associations after the Central Executive network (*r*_adj_ range = -0.0191 to -0.0184 for processing speed; *r*_adj_ range = -0.0274 to -0.0199 for visuospatial ability).

#### **e. Age and cognitive prediction using novel weighting schemes.**

*i. Age prediction.* Across all NOIs, nodes + edges together explained the greatest variance in age (*R*^2^ = 0.223 to 0.537), indicating the collective importance of all elements to brain aging (see Fig. S12). The node*edge and node*edge + node weighting schemes demonstrated approximately similar predictive power to the nodes + edges scheme (*R*^2^ = 0.190 to 0.498 for node*edge; *R*^2^ = 0.221 to 0.501 for node*edge + node), outperforming nodes alone in all NOIs, and outperforming edges alone in all NOIs but PFIT and the whole brain.

*ii. Cognitive prediction.* The contribution of joint weighting schemes to cognitive function varied widely by both network and cognitive domain (see Fig. S14). As with the age prediction results, the three different weighting schemes performed relatively consistently across both domains of cognitive function (*r*’s = -0.193 to 0.099 for nodes + edges; *r*’s = -0.217 to 0.055 for nodes*edges; *r*’s = -0.224 to 0.036 for nodes*edges + nodes). Across all networks, nodes alone outperformed all other weighting schemes (*r*’s = -0.343 to -0.014). Thus, in the context of a LASSO approach that removes redundant predictors of the first-stage outcome (age), information relevant to cognitive function appears to be diluted by the inclusion of edges in addition to nodes.

#### **f. Sensitivity analyses.**

*i. Permuted composites.* Results of the permutation tests are reported in Table S7 and Fig. S9. For edges, the associations between composite scores and processing speed and visuospatial ability were statistically significant relative to the permuted distribution (empirical *p*’s < 0.01). Associations between observed edge-based composites and memory were statistically significant relative to the permuted distribution at empirical *p*’s < 0.05. In contrast, all associations for node composite scores fell within the middle of the permuted distribution for associations with both processing speed and visuospatial ability (empirical *p*’s > 0.08).

*ii. Thresholding analyses.* Both unthresholded age correlations and loadings showed very strong linear relationships with thresholded versions of these metrics (*r*’s = 0.999; Fig. S1). When thresholded edges were fixed to 0, rather than discarded, unthresholded age correlations correlated with thresholded age correlations at *r* = 0.768, and unthresholded loadings correlated with thresholded loadings at *r* = 0.830. These results indicate that potentially spurious connections were unlikely to substantially bias the results reported in this article. Despite this substantial overlap, previous work in this dataset has found that thresholding significantly improves average fractional anisotropy associations with age relative to unthresholded data^1^.

### 3. Supplementary Figures

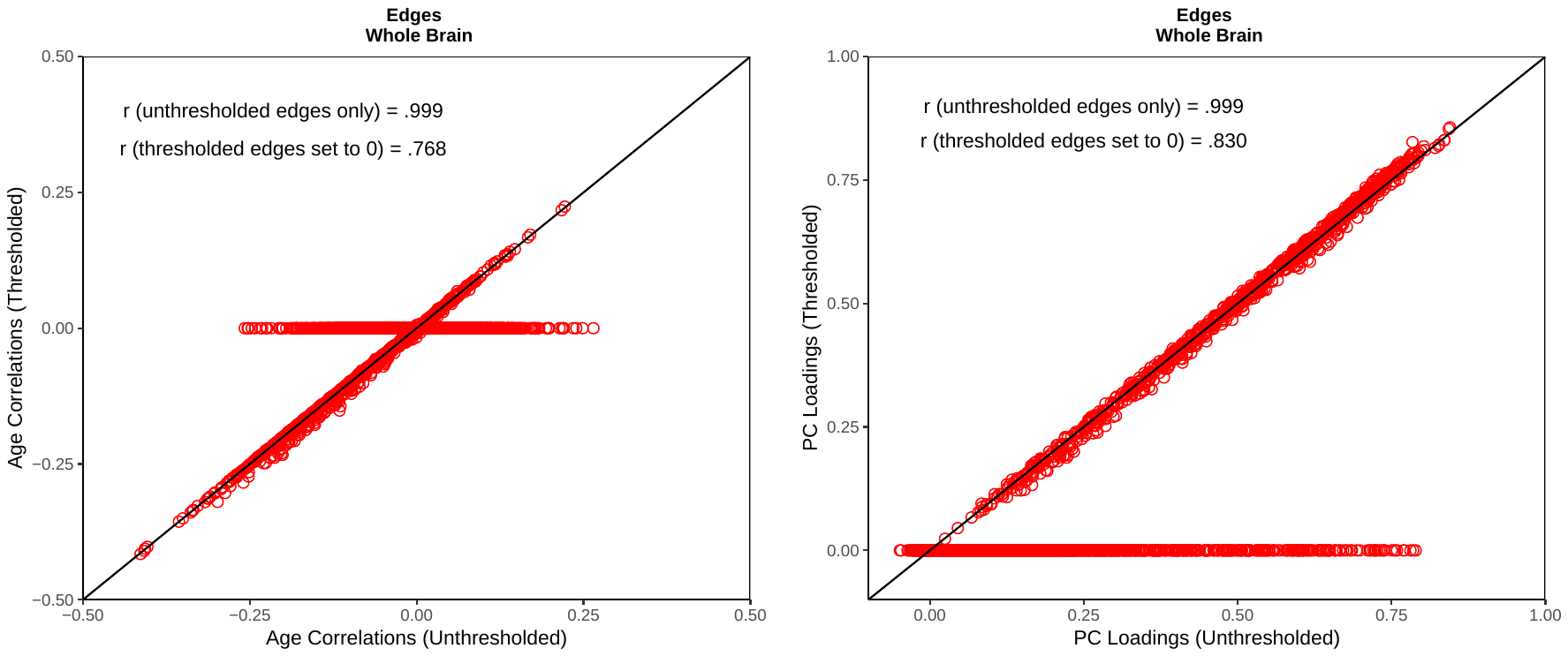

***Figure S1.* Scatterplots of thresholded versus unthresholded edges.** Whole-brain associations between unthresholded and thresholded versions of age correlations and PC loadings. Thresholding was determined by a consistency based-approach^1^. We report correlations both including and excluding thresholded edges (*N* = 2601). Regression line represents a slope of 1 (i.e., X = Y).

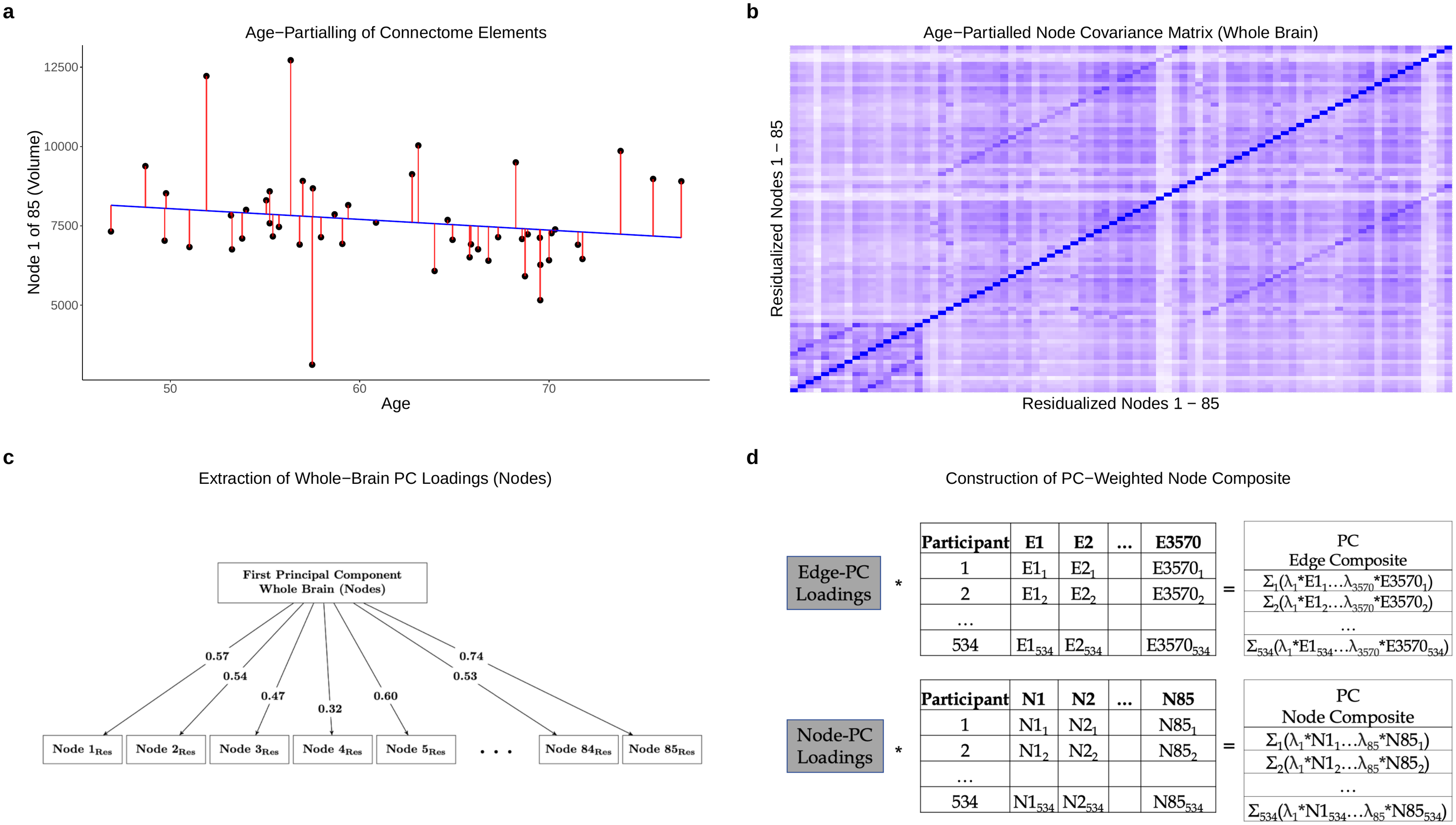

***Figure S2.* Analytic pipeline for obtaining general dimensions of element integrity. a)** Scatterplot between Node 1volume and age in UKB. Red line segments represent each participant’s residual Node 1 integrity after removing age-related variance. This procedure was conducted for every element. Only 50 of the total 3,124 UKB participants are displayed for the sake of visual presentation. **b)** Whole-brain covariance matrix for residualized node integrities. Matrices were subjected to an Eigen decomposition to obtain general dimensions of age-partialled edge and node covariation (i.e., PCs). This procedure was conducted for the 3,564 edges that varied (of 3,570 total) and 85 nodes across the whole brain, as well as within each network, using only the elements contained within each specific NOI. **c)** Extraction of each residualized node’s loading on the first whole-brain PC. We estimated loadings on the first PC derived from the edge and node covariance matrices. **d)** The UKB-derived PC loadings were then used to weight the raw edge and node integrities in LBC1936, such that each LBC1936 participant’s element integrities were multiplied by their respective PC loading and then summed to create a single numerical index of connectome integrity. This procedure was conducted in the whole brain (i.e., using the whole-brain PC loadings) and within each network (i.e., using the network-specific PC loadings and summing only those elements contained within that network), separately for edges and nodes. The same analysis was conducted using the UKB-derived age correlations, instead of PC loadings, as weights.

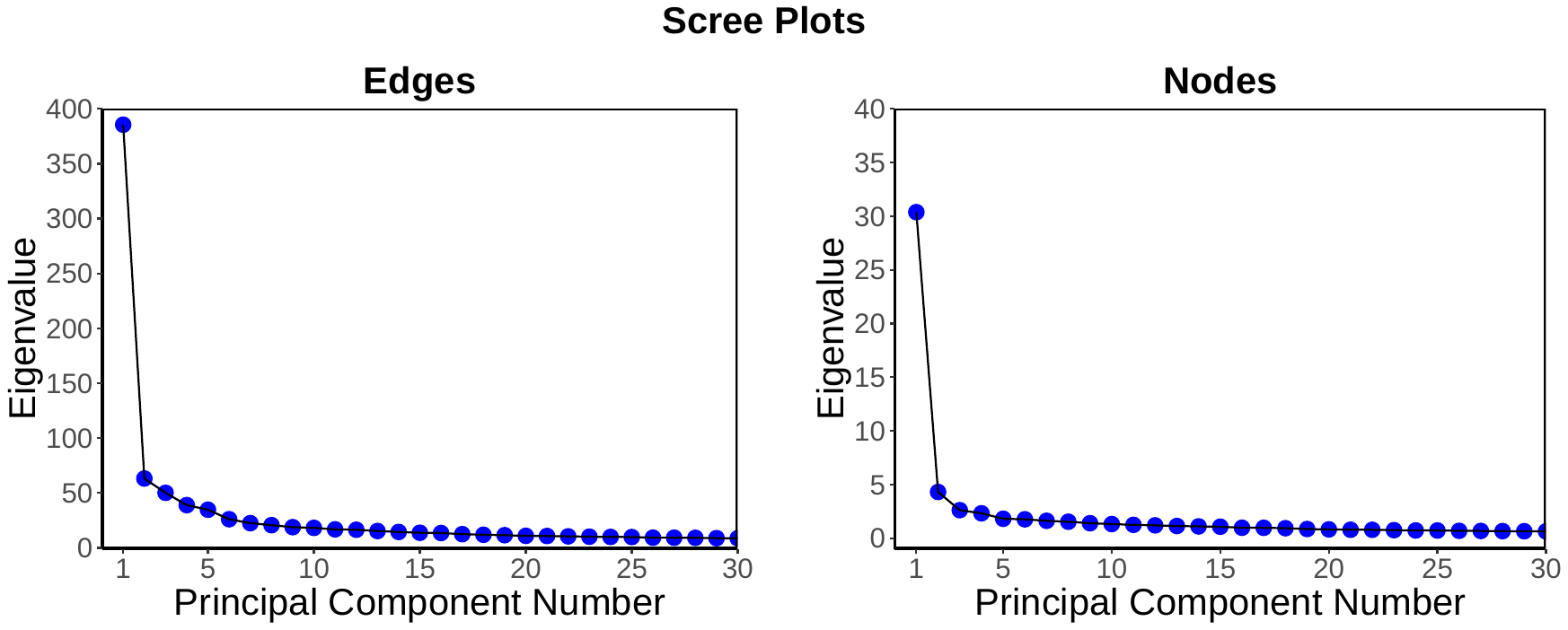

***Figure S3.* Scree plots for whole-brain edge and node Eigen decompositions.** Scree plots reflecting the total variance explained by each principal component for edges and nodes separately. Principal components were estimated from covariance matrices of edges and nodes in UKB participants (*N* = 3,155). Loadings on the first principal component were used to investigate individual differences in global connectome integrity and were used to create weighted connectome integrity composite scores in LBC1936.

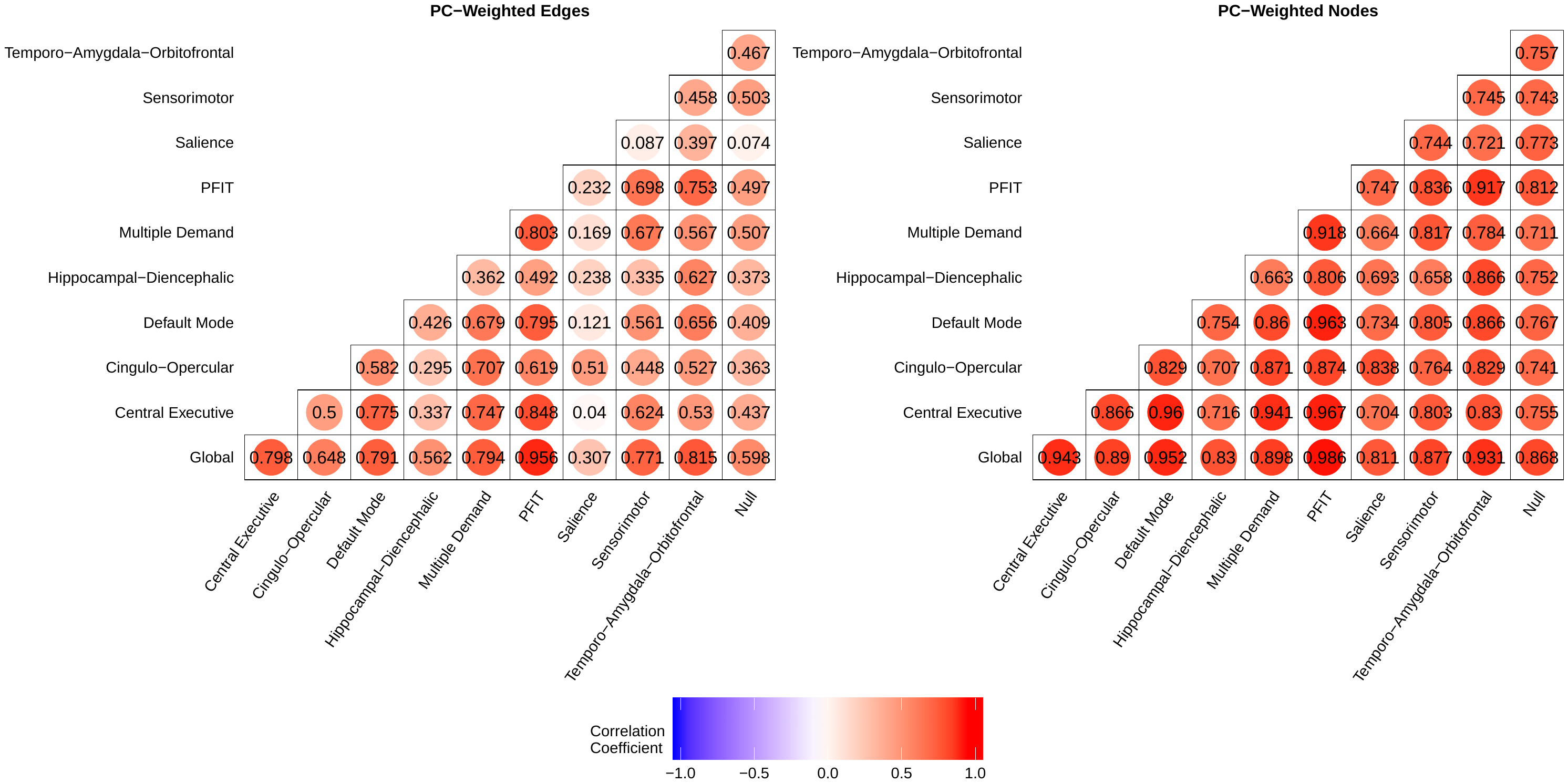

***Figure S4.* Heatmaps of intercorrelations amongst PC-weighted composite scores.** Heatmaps of the correlations between PC-weighted composite scores created in each of the ten prespecified brain networks. Correlations were estimated in edges and nodes separately. The interquartile range for correlations is presented in Table S5. Average correlations between edge- and node-based composites in each network are presented in Table S6.

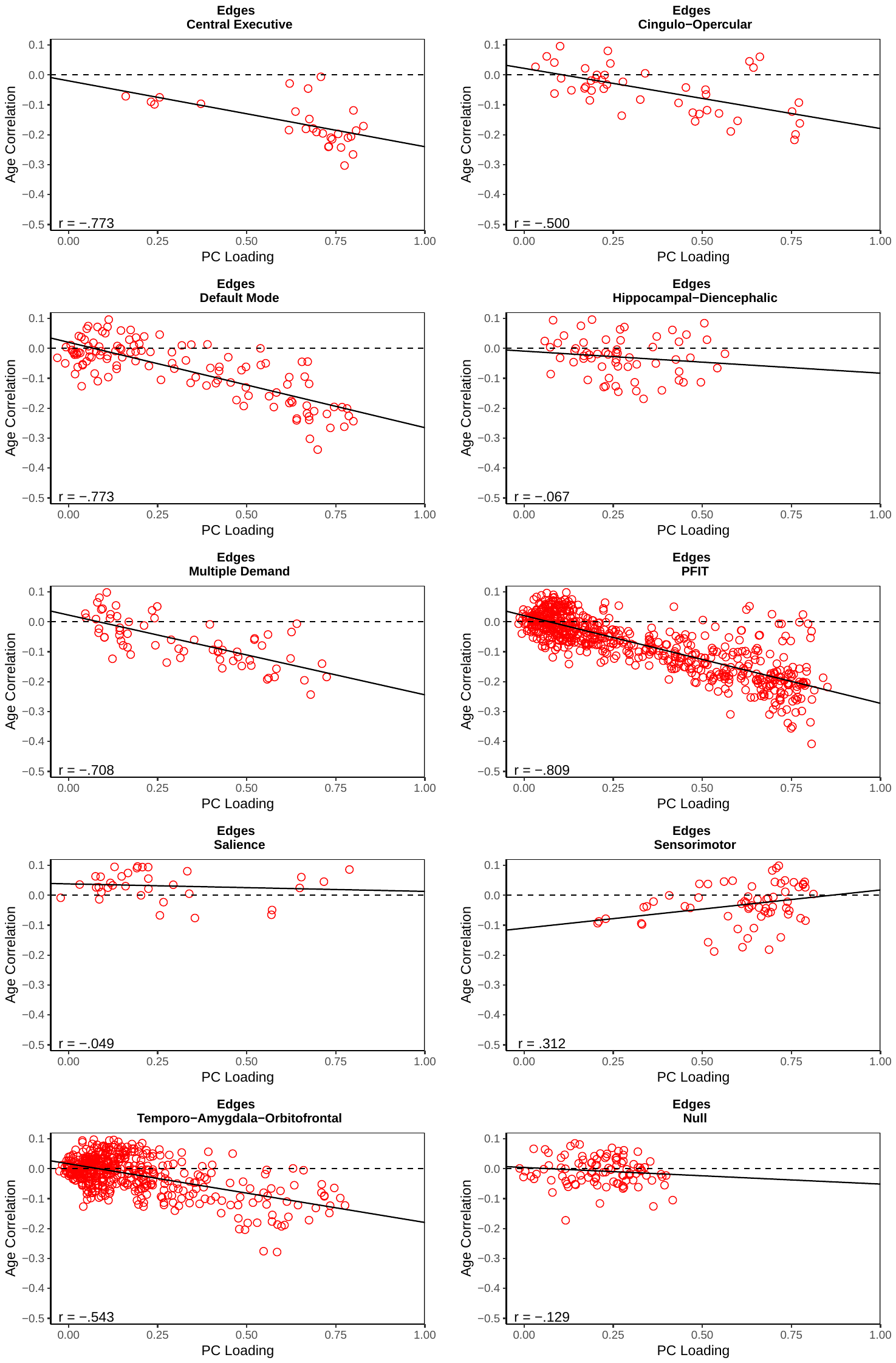

***Figure S5.* Scatterplots of edge-age correlations and PC loadings in NOIs.** Correlations between edge-age correlations and loadings on the first principal component within each NOI. Dashed line represents *r* = 0. Solid line represents the regression line for age correlation on principal component loadings. Principal component loadings were standardized before analysis.

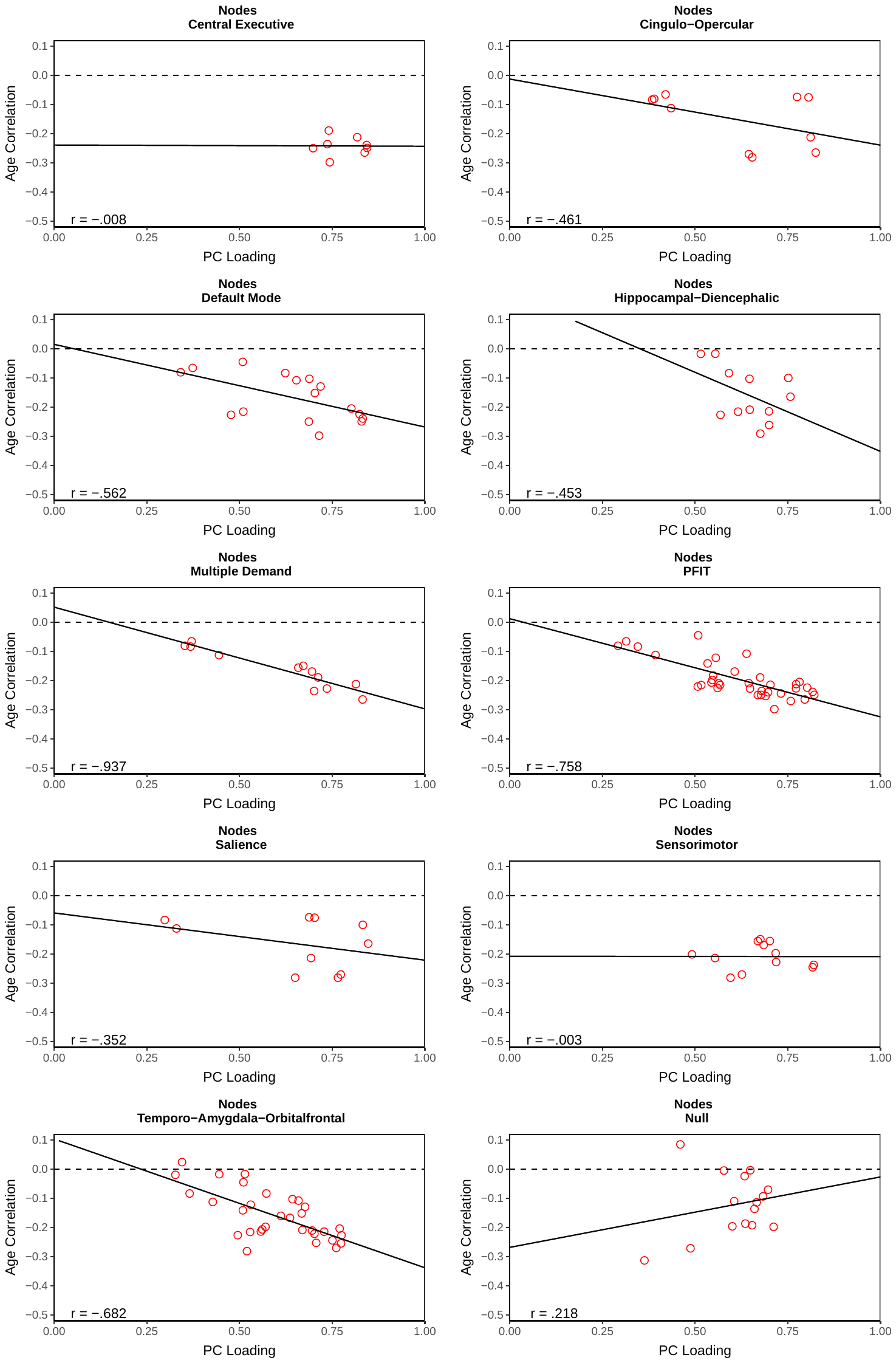

***Figure S6.* Scatterplots of node-age correlations and PC loadings in NOIs**. Correlations between node age correlations and loadings on the first principal component within each NOI. Dashed line represents *r* = 0. Solid line represents the regression line for age correlation on principal component loadings. Principal component loadings were standardized before analysis.

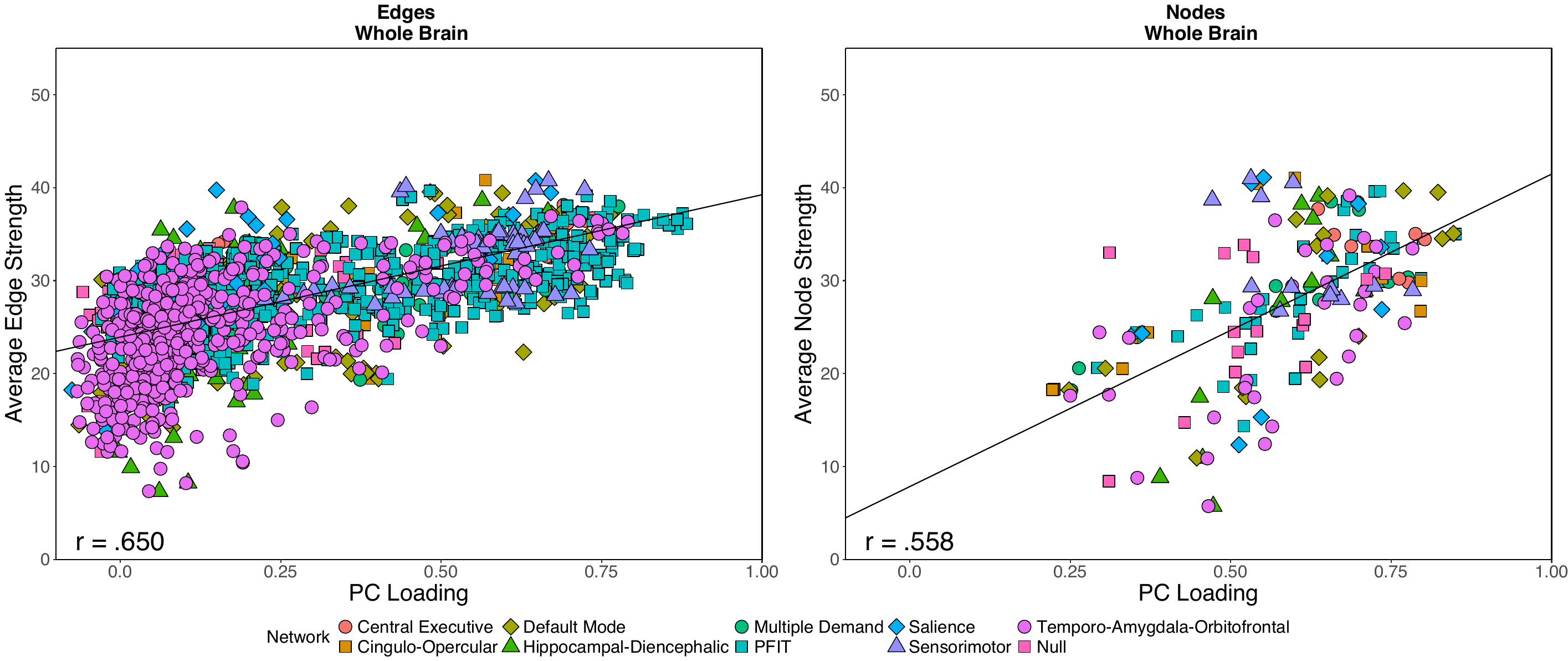

***Figure S7.* Scatterplots of whole-brain topological centrality and PC loadings.** Scatterplots displaying the association between weighted topological centrality (*i.e.,* each connectome element’s average strength) and representativeness of variation in connectome integrity (*i.e.,* each element’s loading on the first principal component). Plots are broken down by element type (*i.e.,* edges and nodes). Each point represents a unique element of the connectome (N=3,564 non-zero edges; 85 nodes). Points are categorized by the network the element belongs to. Line represents the regression line for average strength on principal component loadings.

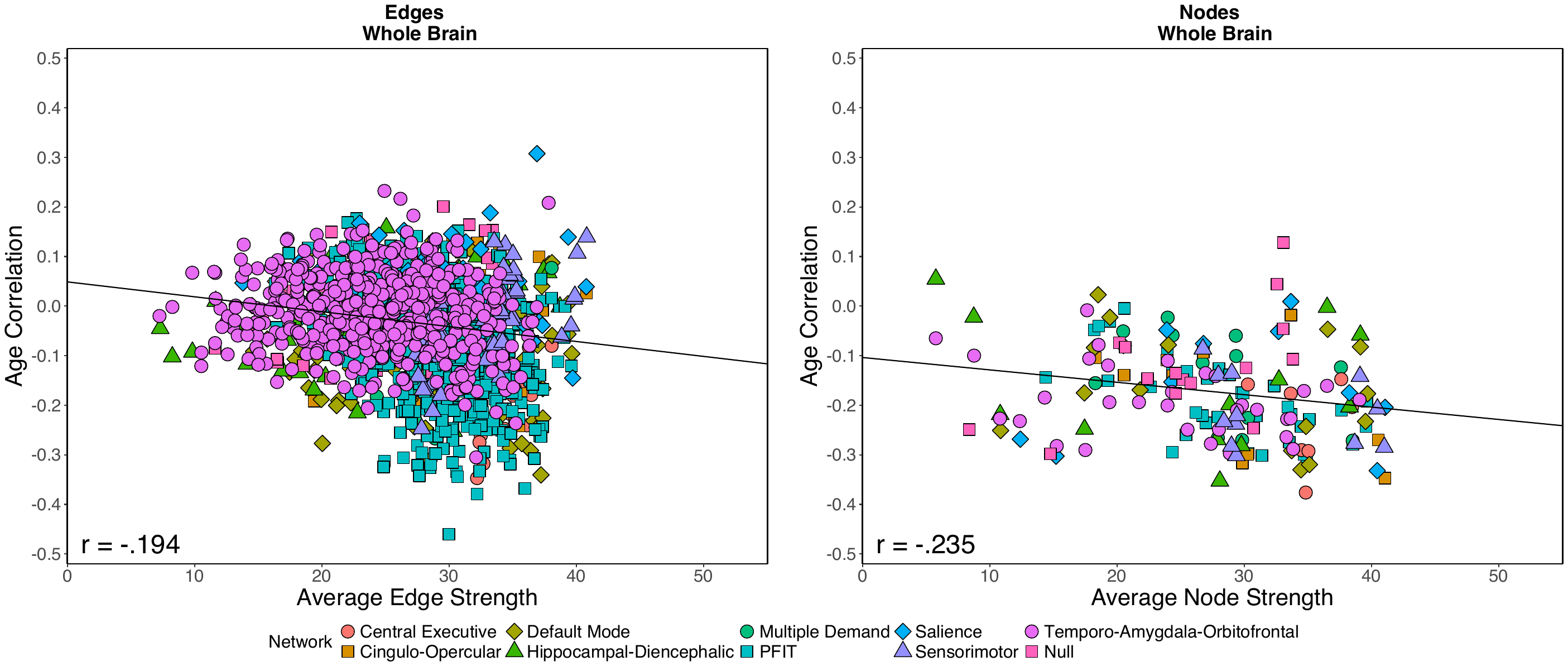

***Figure S8.* Scatterplots of whole-brain topological centrality and age correlations**. Scatterplots displaying the association between each connectome element’s correlation with age and each element’s weighted topological centrality (*i.e.,* each connectome element’s average strength). Plots are broken down by element type (*i.e.,* edges and nodes). Each point represents a unique element of the connectome (N=3,564 non-zero edges; 85 nodes). Points are categorized by the network the element belongs to. Line represents the regression line for each element’s age correlations on average strength.

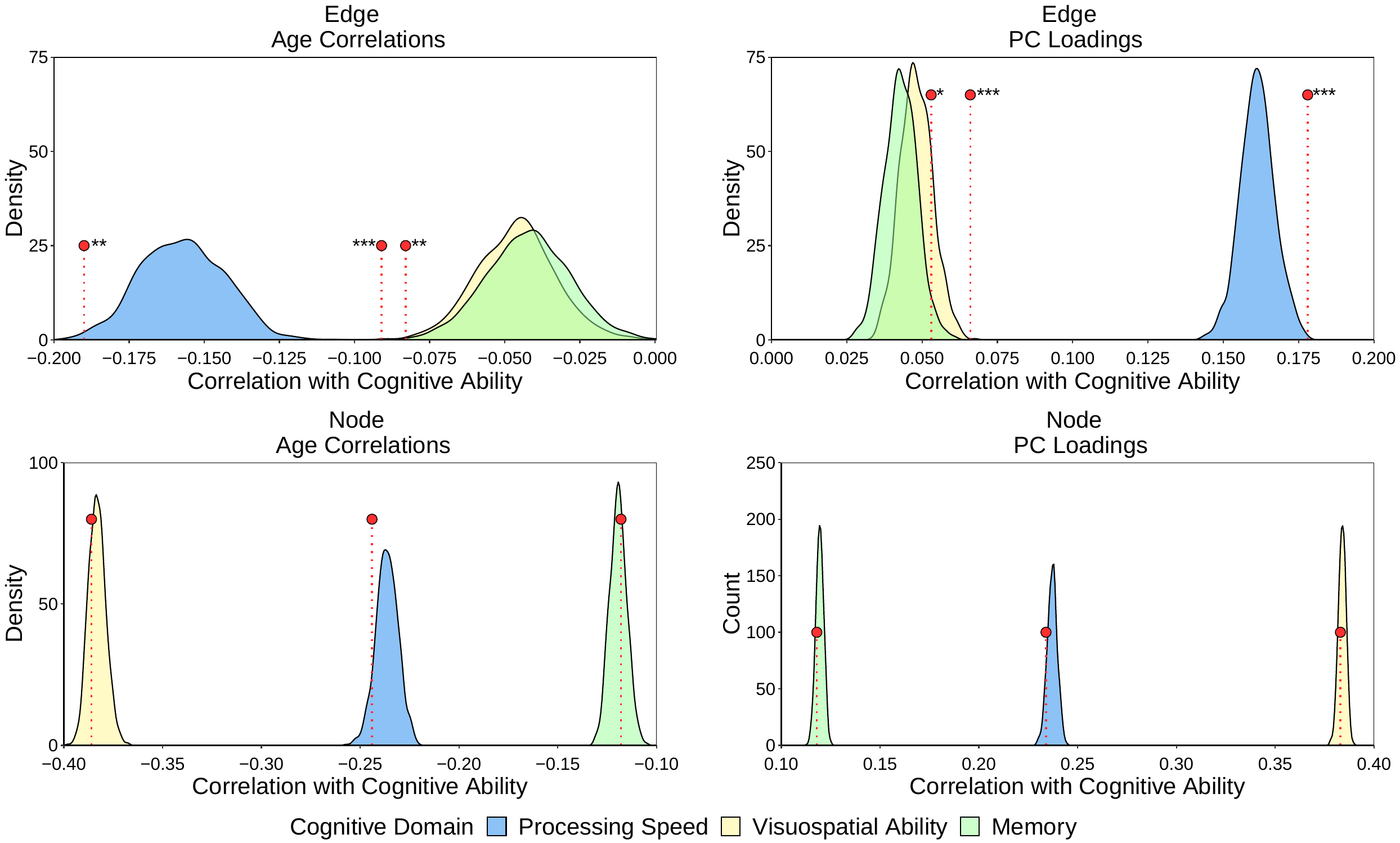

***Figure S9.* Empirical distributions of associations between permuted composite scores and cognitive function.** Empirical distributions of associations between permuted weighted composite scores (*k* = 1000) and domains of cognitive function. Permuted composite scores were created from both age correlations and PC loadings weights in both edges and nodes. Red data points represent observed associations between weighted composite scores and domains of cognitive function (see Fig. 5, Fig. S10, and Table S7 for reference).

* empirical *p* < 0.05; ** empirical *p* < 0.01; *** empirical *p* < 0.001

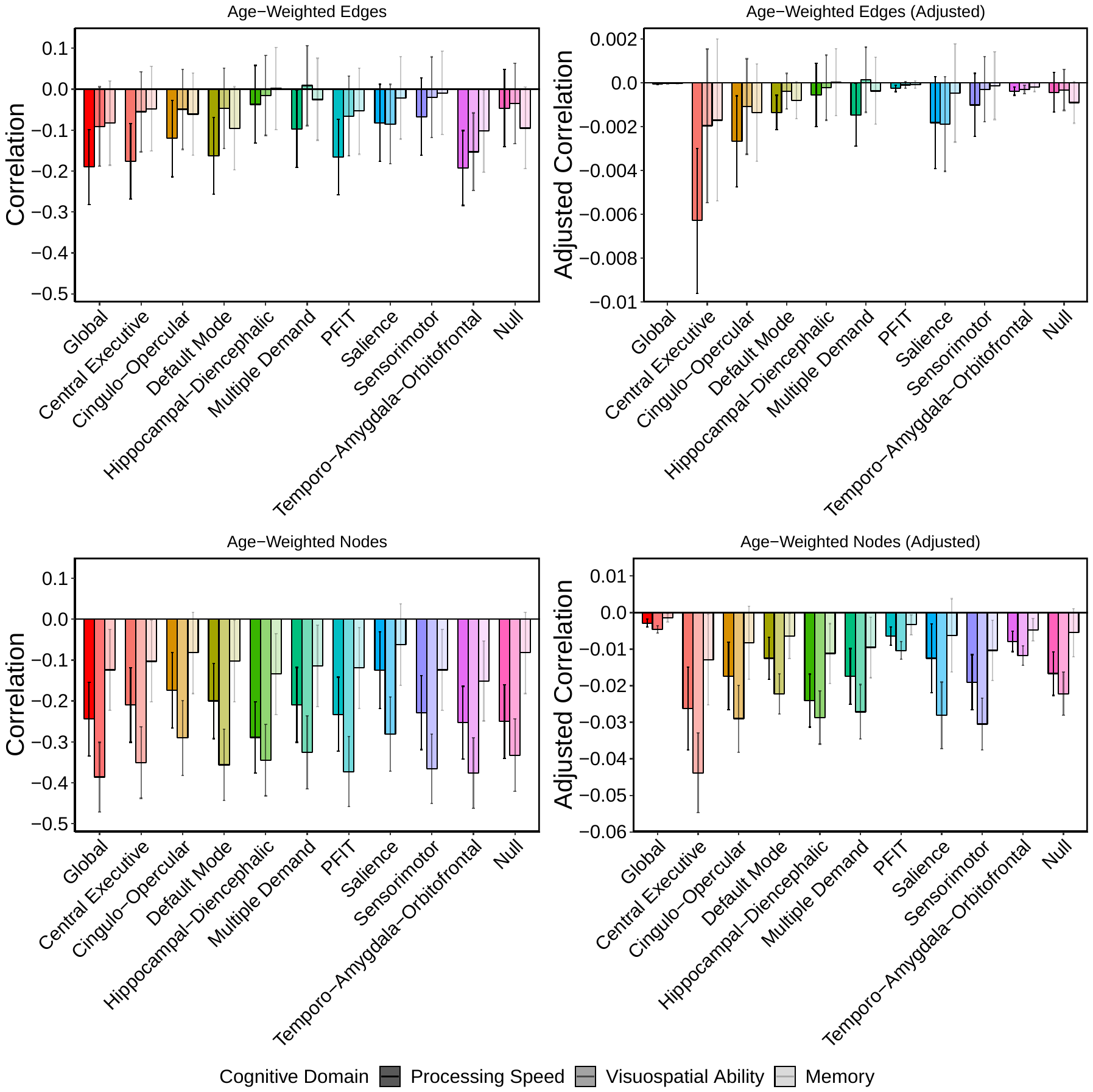

***Figure S10.* Associations between age-weighted composite scores and cognitive function.** Raw and adjusted associations between weighted linear composite scores reflecting age-susceptibility and cognitive function in LBC1936. Scores were created across the whole brain and all NOIs by summing the LBC1936 data weighted by each element’s age correlation discovered in UK Biobank. Plots are broken down by element type (*i.e.*, edges or nodes) and reflect correlations between composite scores from each NOI and the cognitive domains of processing speed, visuospatial ability, and memory.

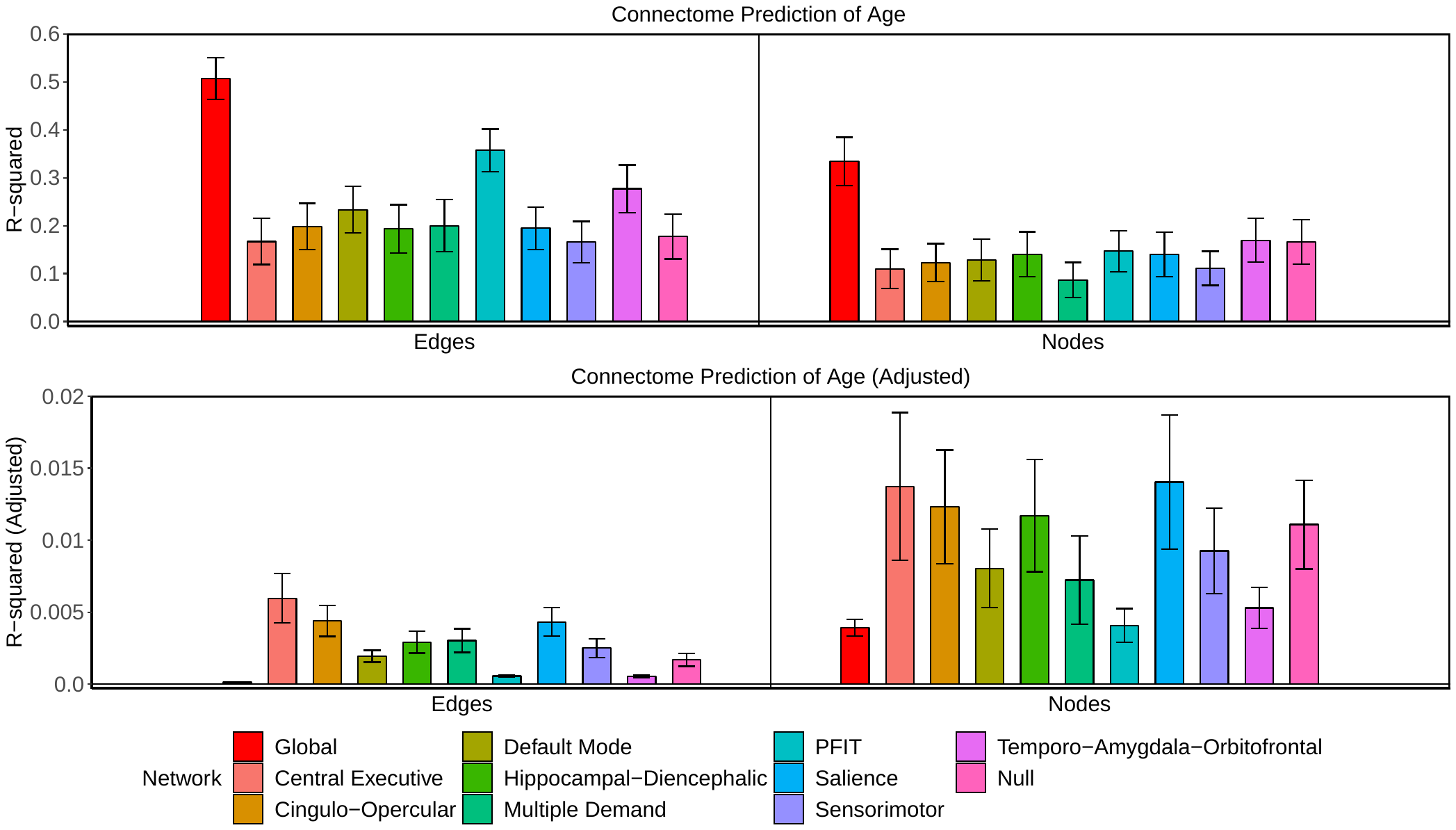

***Figure S11.* LASSO-prediction of age in UKB.** LASSO-prediction of age in a UK Biobank hold-out sample from brain network-specific data trained in UK Biobank. Predictions are broken down by connectome elements (*i.e.*, edges or nodes). Prediction of age is represented as both the raw and adjusted *R^2^* value, with error bars representing bootstrapped 95% confidence intervals based on the 100 iterations of *R^2^* calculation. A full description of these findings can be found in the Supplementary Results.

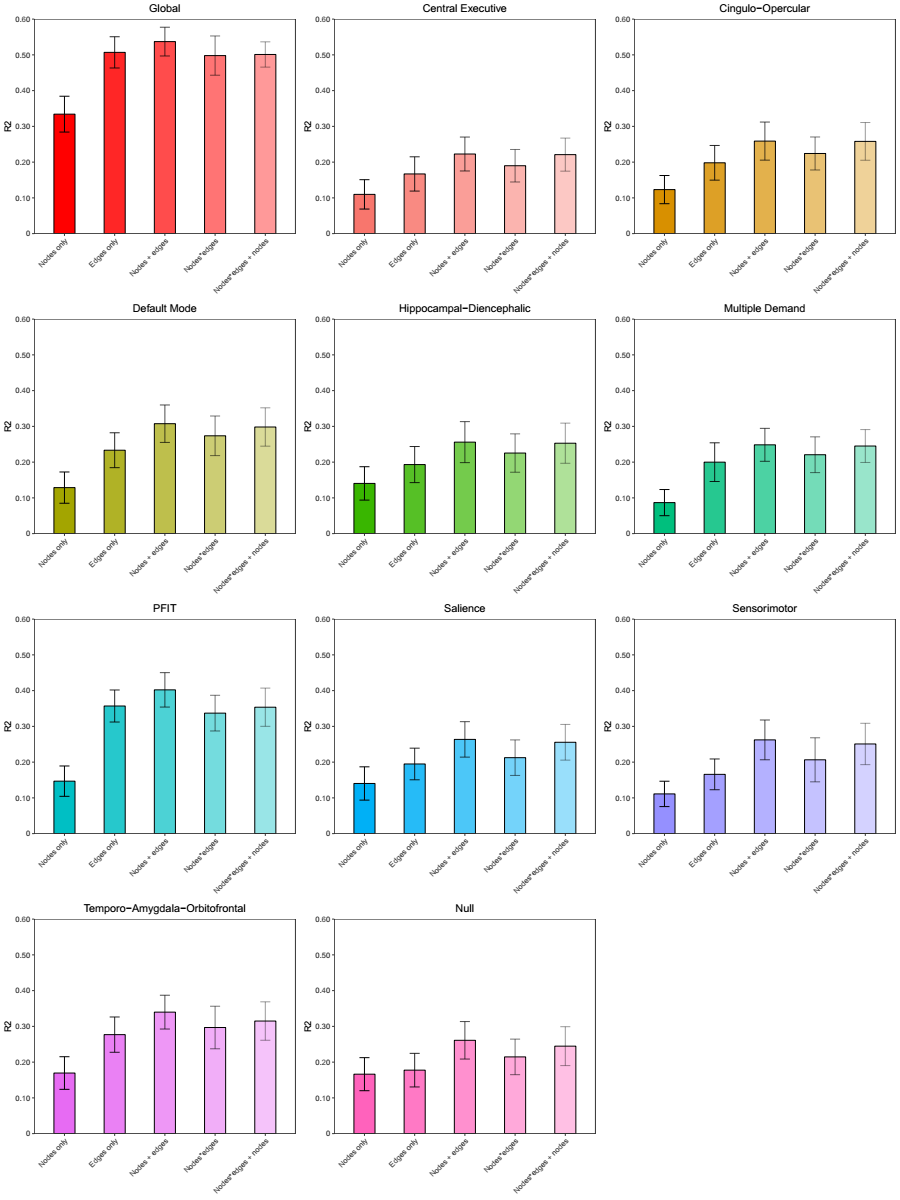
 ***Figure S12.* LASSO-prediction of age in UKB using novel weighting schemes.** LASSO-prediction of age in each NOI broken down by weighting scheme (nodes alone, edges alone, nodes + edges, nodes*edges, nodes*edges + nodes). A full description of each weighting scheme can be found in the Supplementary Method.

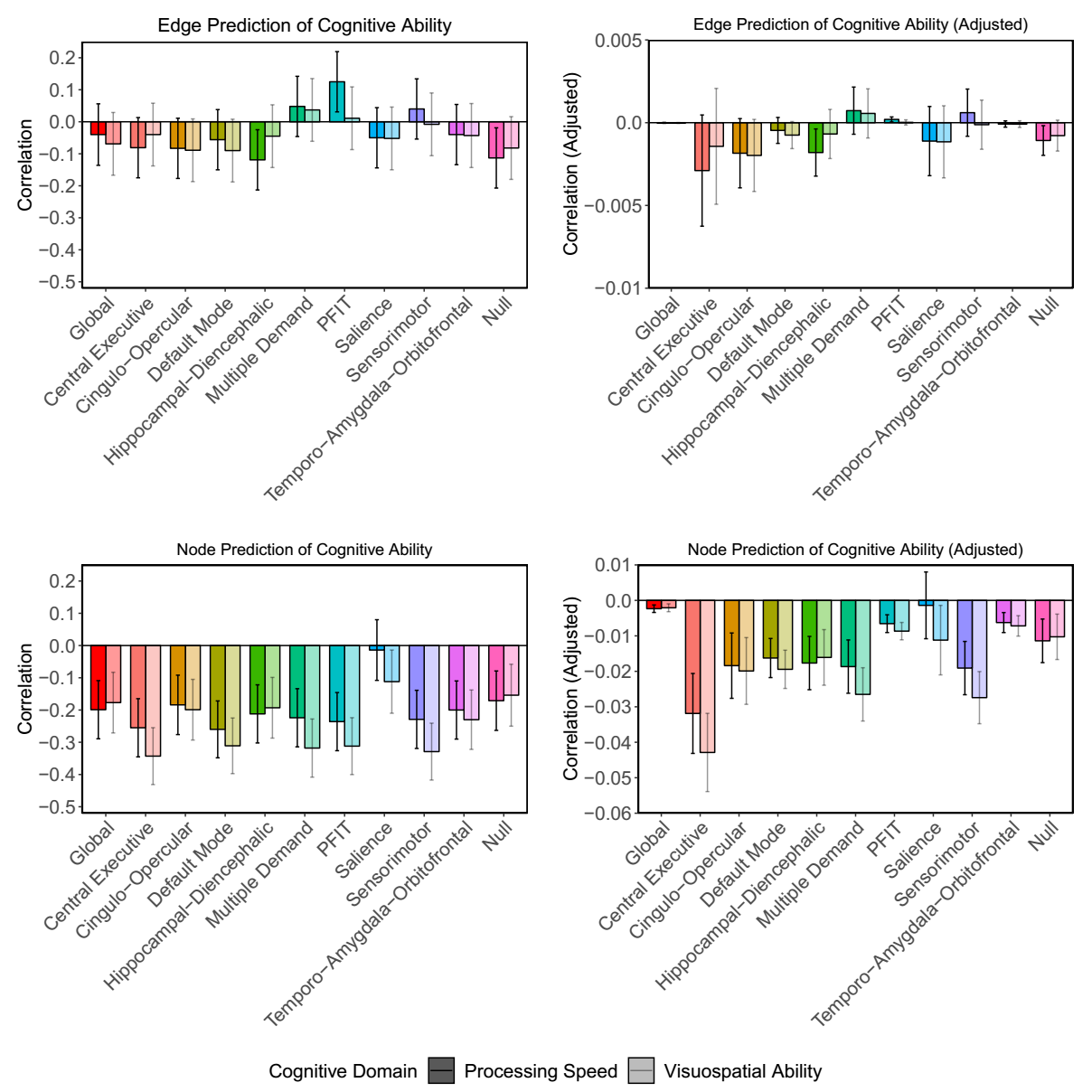

***Figure S13.* LASSO-prediction of cognitive function in LBC1936.** LASSO-prediction of cognitive function (processing speed and visuospatial ability) in LBC1936 from brain network-specific age data trained in UK Biobank. Predictions are broken down by connectome elements (*i.e.*, edges or nodes). Prediction of cognitive function is represented as both the raw and adjusted correlation between LASSO-retained element-age coefficients from each NOI and latent factors of processing speed and visuospatial ability, with error bars representing 95% confidence intervals. A full description of these results can be found in the Supplementary Results.

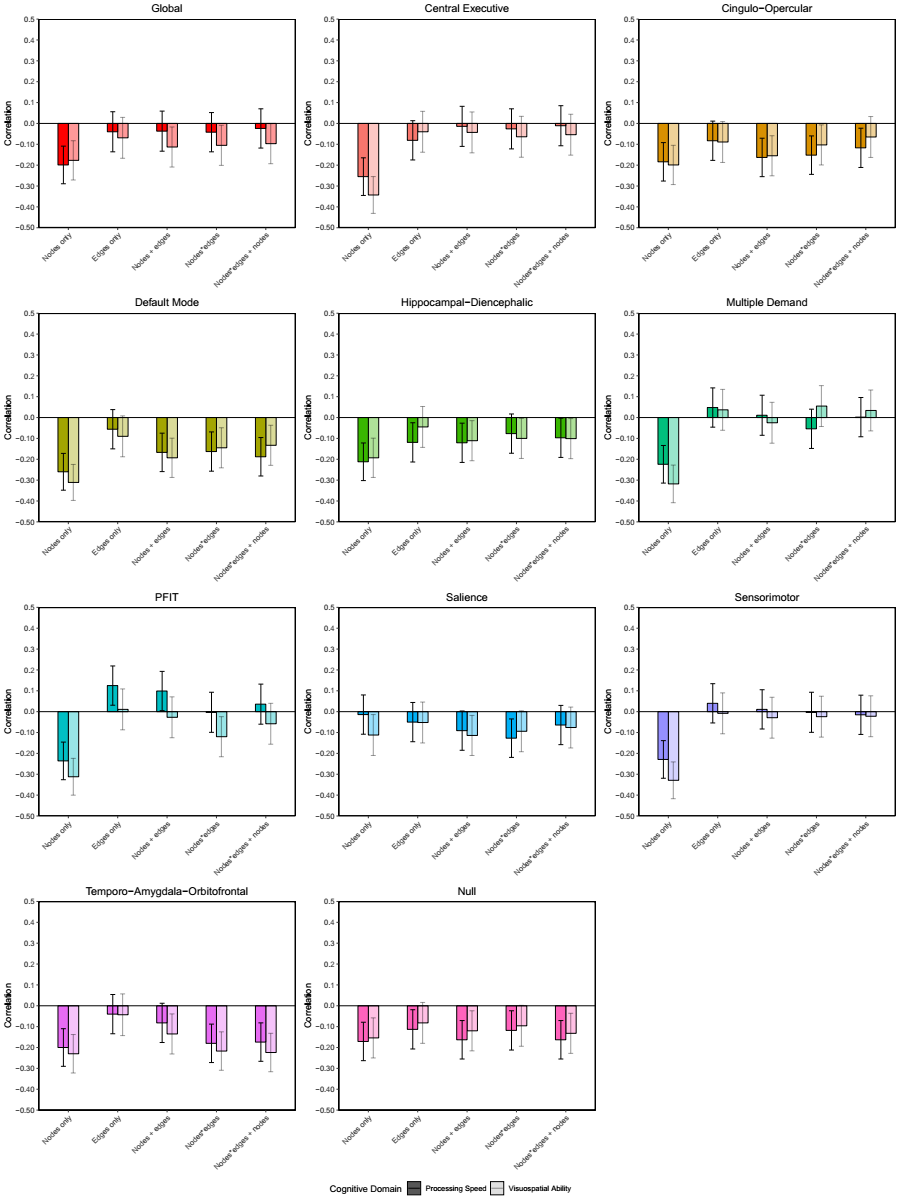

***Figure S14.* LASSO-prediction of cognitive function in LBC1936 using novel weighting schemes.** LASSO-prediction of cognitive function in each network-of-interest broken down by weighting scheme (nodes alone, edges alone, nodes + edges, nodes*edges, nodes*edges + nodes). A full description of each weighting scheme can be found in the Supplementary Method.

### 4. Supplementary Tables

#### ***Table S1.* Properties of each brain NOI.** Properties of each brain NOI, with a canonical reference describing the network’s makeup, previous associations and elements. See Fig. 1 for illustration of network properties.

| Network | Number of Nodes | Number of Edges | Hypothesis | Select Regions | Implicated in |
| --- | --- | --- | --- | --- | --- |
| All networks (global) | 85 | 3570 | + |  |  |
| P-FIT^2^ | 36 | 630 | + | DLPFC, inferior and superior parietal lobule, anterior cingulate, and specific regions within the temporal and occipital lobes | General intelligence |
| Central Executive^3,4,5^ | 8 | 28 | + | rDLPFC, posterior parietal cortex | Activation associated with selecting, switching, and attending to salient events |
| Multiple Demand^6,7^ | 12 | 66 | + | middle frontal, inferior parietal, pre-SMA, anterior cingulate, rostral prefrontal, insula/frontal operculum | General purpose activation in cognitive demanding tasks, suggesting a role in cognitive flexibility, executive control, and abstract problem solving |
| Cingulo-Opercular^5^ | 10 | 45 | + | dorsal anterior cingulate, superior and anterior frontal cortex, insula, thalamus | Stable set control, maintenance of task-relevant sustained attention |
| Default Mode^8,9^ | 16 | 120 | - | ventromedial frontal, medial temporal and posterior cingulate cortices, angular gyrus and cingulum bundle | Extensive de-activation in functional MRI during cognitively demanding tasks |
| Hippocampal-Diencephalic^10,11^ | 12 | 66 | + | hippocampus, diencephalon, ventral cingulum and fornix | Memory and spatial orientation |
| Salience^4,12^ | 10 | 45 | - | insula, anterior cingulate cortex, amydala, substantia nigra/VTA and thalamus | Orientation of attention to the most homeostatically relevant even from moment to moment |
| Sensorimotor^13,14^ | 12 | 66 | - | precentral, postcentral, pre and post SMA, caudal cingulate, caudal middle frontal, thalamus, putamen | Initiation and control of movements |
| Temporo-Amygdala-Orbital^9,15^ | 32 | 496 | - | anterior temporal cortex, amygdala and orbitofrontal cortices, ACC, and parts of cingulum bundle | Visceral emotion and sensation |
| Null | 15 | 105 | - | ROIs not included in any of the other subnetworks | - |
| *Note.* For each network, the number of edges is N*((N-1)/2) the number of nodes. + refers to subnetwork for which we hypothesized a positive association between subnetwork integrity and cognitive function. – refers to a negative control network, i.e. a network for which we do not hypothesize a positive association between subnetwork integrity and cognitive function. | | | | | |

***Table S2.* Assignments of each brain region to each of the ten NOIs.** Brain regions were parceled per the Desikan-Killiany atlas.

| *Node* | *Central Executive* | *Cingulo-Opercular* | *Default Mode* | *Hippocampal-Diencephalic* | *Multiple Demand* | *PFIT* | *Salience* | *Sensori-motor* | *Temporo-Amygdala-Orbitofrontal* | *Null* |
| --- | --- | --- | --- | --- | --- | --- | --- | --- | --- | --- |
| Left-thalamus | 0 | 1 | 0 | 0 | 0 | 0 | 1 | 1 | 0 | 0 |
| Left-caudate | 0 | 0 | 0 | 0 | 0 | 0 | 0 | 0 | 0 | 1 |
| Left-putamen | 0 | 0 | 0 | 0 | 0 | 0 | 0 | 1 | 0 | 0 |
| Left-pallidum | 0 | 0 | 0 | 0 | 0 | 0 | 0 | 0 | 0 | 1 |
| Brain stem | 0 | 0 | 0 | 0 | 0 | 0 | 0 | 0 | 0 | 1 |
| Left-hippocampus | 0 | 0 | 0 | 1 | 0 | 0 | 0 | 0 | 0 | 0 |
| Left-amygdala | 0 | 0 | 0 | 0 | 0 | 0 | 1 | 0 | 1 | 0 |
| Left-accumbens area | 0 | 0 | 0 | 0 | 0 | 0 | 0 | 0 | 0 | 1 |
| Left-ventral diencephalon | 0 | 0 | 0 | 1 | 0 | 0 | 1 | 0 | 0 | 0 |
| Right-thalamus | 0 | 1 | 0 | 0 | 0 | 0 | 1 | 1 | 0 | 0 |
| Right-caudate | 0 | 0 | 0 | 0 | 0 | 0 | 0 | 0 | 0 | 1 |
| Right-putamen | 0 | 0 | 0 | 0 | 0 | 0 | 0 | 1 | 0 | 0 |
| Right-pallidum | 0 | 0 | 0 | 0 | 0 | 0 | 0 | 0 | 0 | 1 |
| Right-hippocampus | 0 | 0 | 0 | 1 | 0 | 0 | 0 | 0 | 0 | 0 |
| Right-amygdala | 0 | 0 | 0 | 0 | 0 | 0 | 1 | 0 | 1 | 0 |
| Right-accumbens area | 0 | 0 | 0 | 0 | 0 | 0 | 0 | 0 | 0 | 1 |
| Right-ventral diencephalon | 0 | 0 | 0 | 1 | 0 | 0 | 1 | 0 | 0 | 0 |
| Left-superior temporal sulcus | 0 | 0 | 0 | 0 | 0 | 1 | 0 | 0 | 1 | 0 |
| Left-caudal anterior cingulate | 0 | 1 | 0 | 0 | 1 | 1 | 1 | 0 | 1 | 0 |
| Left-caudal middle frontal | 0 | 0 | 0 | 0 | 1 | 1 | 0 | 1 | 0 | 0 |
| Left-cuneus | 0 | 0 | 0 | 0 | 0 | 0 | 0 | 0 | 0 | 1 |
| Left-entorhinal | 0 | 0 | 0 | 1 | 0 | 0 | 0 | 0 | 1 | 0 |
| Left-fusiform | 0 | 0 | 0 | 1 | 0 | 1 | 0 | 0 | 1 | 0 |
| Left-inferior parietal | 1 | 0 | 1 | 0 | 0 | 1 | 0 | 0 | 0 | 0 |
| Left-inferior temporal | 0 | 0 | 0 | 0 | 0 | 0 | 0 | 0 | 1 | 0 |
| Left-isthmus cingulate | 0 | 0 | 1 | 1 | 0 | 0 | 0 | 0 | 1 | 0 |
| Left-lateral occipital | 0 | 0 | 0 | 0 | 0 | 1 | 0 | 0 | 0 | 0 |
| Left-lateral orbitofrontal | 0 | 0 | 0 | 0 | 0 | 0 | 0 | 0 | 1 | 0 |
| Left-lingual | 0 | 0 | 0 | 0 | 0 | 0 | 0 | 0 | 0 | 1 |
| Left-medial orbitofrontal | 0 | 0 | 1 | 0 | 0 | 0 | 0 | 0 | 1 | 0 |
| Left-middle temporal | 0 | 0 | 0 | 0 | 0 | 1 | 0 | 0 | 1 | 0 |
| Left-parahippocampal | 0 | 0 | 1 | 1 | 0 | 0 | 0 | 0 | 1 | 0 |
| Left-paracentral | 0 | 0 | 0 | 0 | 1 | 0 | 0 | 1 | 0 | 0 |
| Left-pars opercularis | 0 | 0 | 0 | 0 | 0 | 1 | 0 | 0 | 0 | 0 |
| Left-pars orbitalis | 0 | 0 | 0 | 0 | 0 | 1 | 0 | 0 | 0 | 0 |

***Table S2, continued***

| *Node* | *Central Executive* | *Cingulo-Opercular* | *Default Mode* | *Hippocampal-Diencephalic* | *Multiple Demand* | *PFIT* | *Salience* | *Sensori-motor* | *Temporo-Amygdala-Orbitofrontal* | *Null* |
| --- | --- | --- | --- | --- | --- | --- | --- | --- | --- | --- |
| Left-pars triangularis | 0 | 0 | 0 | 0 | 0 | 1 | 0 | 0 | 0 | 0 |
| Left-pericalcarine | 0 | 0 | 0 | 0 | 0 | 0 | 0 | 0 | 0 | 1 |
| Left-postcentral | 0 | 0 | 0 | 0 | 0 | 0 | 0 | 1 | 0 | 0 |
| Left-posterior cingulate | 0 | 0 | 0 | 0 | 0 | 0 | 0 | 0 | 1 | 0 |
| Left-precentral | 0 | 0 | 0 | 0 | 0 | 0 | 0 | 1 | 0 | 0 |
| Left-precuneus | 0 | 0 | 1 | 0 | 0 | 1 | 0 | 0 | 0 | 0 |
| Left-rostral anterior cingulate | 0 | 0 | 1 | 0 | 0 | 1 | 0 | 0 | 1 | 0 |
| Left-rostral middle frontal | 1 | 1 | 0 | 0 | 1 | 1 | 0 | 0 | 0 | 0 |
| Left-superior frontal | 1 | 0 | 1 | 0 | 0 | 1 | 0 | 0 | 0 | 0 |
| Left-superior parietal | 1 | 0 | 0 | 0 | 1 | 1 | 0 | 0 | 0 | 0 |
| Left-superior temporal | 0 | 0 | 0 | 0 | 0 | 1 | 0 | 0 | 1 | 0 |
| Left-supramarginal | 0 | 0 | 0 | 0 | 0 | 0 | 0 | 0 | 0 | 1 |
| Left-frontal pole | 0 | 1 | 1 | 0 | 1 | 1 | 0 | 0 | 0 | 0 |
| Left-temporal pole | 0 | 0 | 0 | 0 | 0 | 0 | 0 | 0 | 1 | 0 |
| Left-transverse temporal | 0 | 0 | 0 | 0 | 0 | 1 | 0 | 0 | 1 | 0 |
| Left-insula | 0 | 1 | 0 | 0 | 0 | 0 | 1 | 0 | 0 | 0 |
| Right-superior temporal sulcus | 0 | 0 | 0 | 0 | 0 | 1 | 0 | 0 | 1 | 0 |
| Right-caudal anterior cingulate | 0 | 1 | 0 | 0 | 1 | 1 | 1 | 0 | 1 | 0 |
| Right-caudal middle frontal | 0 | 0 | 0 | 0 | 1 | 1 | 0 | 1 | 0 | 0 |
| Right-cuneus | 0 | 0 | 0 | 0 | 0 | 0 | 0 | 0 | 0 | 1 |
| Right-entorhinal | 0 | 0 | 0 | 1 | 0 | 0 | 0 | 0 | 1 | 0 |
| Right-fusiform | 0 | 0 | 0 | 1 | 0 | 1 | 0 | 0 | 1 | 0 |
| Right-inferior parietal | 1 | 0 | 1 | 0 | 0 | 1 | 0 | 0 | 0 | 0 |
| Right-inferior temporal | 0 | 0 | 0 | 0 | 0 | 0 | 0 | 0 | 1 | 0 |
| Right-isthmus cingulate | 0 | 0 | 1 | 1 | 0 | 0 | 0 | 0 | 1 | 0 |
| Right-lateral occipital | 0 | 0 | 0 | 0 | 0 | 1 | 0 | 0 | 0 | 0 |
| Right-lateral orbitofrontal | 0 | 0 | 0 | 0 | 0 | 0 | 0 | 0 | 1 | 0 |
| Right-lingual | 0 | 0 | 0 | 0 | 0 | 0 | 0 | 0 | 0 | 1 |
| Right-medial orbitofrontal | 0 | 0 | 1 | 0 | 0 | 0 | 0 | 0 | 1 | 0 |
| Right-middle temporal | 0 | 0 | 0 | 0 | 0 | 1 | 0 | 0 | 1 | 0 |
| Right-parahippocampal | 0 | 0 | 1 | 1 | 0 | 0 | 0 | 0 | 1 | 0 |
| Right-paracentral | 0 | 0 | 0 | 0 | 1 | 0 | 0 | 1 | 0 | 0 |
| Right-pars opercularis | 0 | 0 | 0 | 0 | 0 | 1 | 0 | 0 | 0 | 0 |
| Right-pars orbitalis | 0 | 0 | 0 | 0 | 0 | 1 | 0 | 0 | 0 | 0 |
| Right-pars triangularis | 0 | 0 | 0 | 0 | 0 | 1 | 0 | 0 | 0 | 0 |

***Table S2, continued***

| *Node* | *Central Executive* | *Cingulo-Opercular* | *Default Mode* | *Hippocampal-Diencephalic* | *Multiple Demand* | *PFIT* | *Salience* | *Sensori-motor* | *Temporo-Amygdala-Orbitofrontal* | *Null* |
| --- | --- | --- | --- | --- | --- | --- | --- | --- | --- | --- |
| Right-pericalcarine | 0 | 0 | 0 | 0 | 0 | 0 | 0 | 0 | 0 | 1 |
| Right-postcentral | 0 | 0 | 0 | 0 | 0 | 0 | 0 | 1 | 0 | 0 |
| Right-posterior cingulate | 0 | 0 | 0 | 0 | 0 | 0 | 0 | 0 | 1 | 0 |
| Right-precentral | 0 | 0 | 0 | 0 | 0 | 0 | 0 | 1 | 0 | 0 |
| Right-precuneus | 0 | 0 | 1 | 0 | 0 | 1 | 0 | 0 | 0 | 0 |
| Right-rostral anterior cingulate | 0 | 0 | 1 | 0 | 0 | 1 | 0 | 0 | 1 | 0 |
| Right-rostral middle frontal | 1 | 1 | 0 | 0 | 1 | 1 | 0 | 0 | 0 | 0 |
| Right-superior frontal | 1 | 0 | 1 | 0 | 0 | 1 | 0 | 0 | 0 | 0 |
| Right-superior parietal | 1 | 0 | 0 | 0 | 1 | 1 | 0 | 0 | 0 | 0 |
| Right-superior temporal | 0 | 0 | 0 | 0 | 0 | 1 | 0 | 0 | 1 | 0 |
| Right-supramarginal | 0 | 0 | 0 | 0 | 0 | 0 | 0 | 0 | 0 | 1 |
| Right-frontal pole | 0 | 1 | 1 | 0 | 1 | 1 | 0 | 0 | 0 | 0 |
| Right-temporal pole | 0 | 0 | 0 | 0 | 0 | 0 | 0 | 0 | 1 | 0 |
| Right-transverse temporal | 0 | 0 | 0 | 0 | 0 | 1 | 0 | 0 | 1 | 0 |
| Right-insula | 0 | 1 | 0 | 0 | 0 | 0 | 1 | 0 | 0 | 0 |

#### ***Table S3.* Unique elements within each NOI.**

| **Network** | **Total Nodes** | **Unique Nodes** | **Total Edges** | **Unique Edges** |
| --- | --- | --- | --- | --- |
| Central Executive | 8 | 0 | 28 | 0 |
| Cingulo-Opercular | 10 | 0 | 45 | 16 |
| Default Mode | 16 | 0 | 120 | 48 |
| Hippocampal-Diencephalic | 12 | 2 | 66 | 37 |
| Multiple Demand | 12 | 0 | 66 | 16 |
| PFIT | 36 | 8 | 630 | 436 |
| Salience | 10 | 0 | 45 | 24 |
| Sensorimotor | 12 | 6 | 66 | 59 |
| Temporo-Amygdala-Orbitofrontal | 32 | 8 | 496 | 352 |
| Null | 15 | 15 | 105 | 105 |

*Note.* Within each network, the total number of edges is equal to N*((N-1)/2), where N equals the number of nodes. This same mathematical principal does not apply to the estimation of unique elements within a network. Two nodes may be present across several networks, but may only be jointly present within a single subnetwork, thus returning a unique edge for that subnetwork, but not unique nodes. 39 out of 85 nodes are unique specific networks, though note that 40% of these unique nodes are part of the Null network, which by definition contains elements not a part of any other network. 2210 edges are not a part of any network because they connect two nodes that are not jointly present within a single subnetwork. Of the edges that occur within networks, 1093 out 1360 are unique.

#### ***Table S4.*** **Average age-element associations within NOIs.**

| **Network** | **Average Age-Edge Correlation** | **Average Age-Node Correlation** |
| --- | --- | --- |
| Global | -.034 (.086) | -.172 (.084) |
| Central Executive | -.161 (.075) | -.242 (.033) |
| Cingulo-Opercular | -.048 (.081) | -.152 (.093) |
| Default Mode | -.066 (.095) | -.167 (.080) |
| Hippocampal-Diencephalic | -.017 (.077) | -.158 (.092) |
| Multiple Demand | -.064 (.076) | -.162 (.066) |
| PFIT | -.070 (.094) | -.197 (.062) |
| Salience | .061 (.071) | -.166 (.088) |
| Sensorimotor | -.027 (.069) | -.209 (.045) |
| Temporo-Amygdala-Orbitofrontal | -.012 (.063) | -.159 (.083) |
| Null | -.0005 (.056) | -.122 (.108) |

*Note.* Standard deviations are in parentheses.

***Table S5.* Interquartile range and Eigen decomposition for PC-weighted composite scores.** Interquartile range and percentage of variation explained by first principal component for correlations between PC-weighted linear composites across the ten NOIs. Ranges include composites created for the whole brain and null networks.

| ***Element*** | ***0%*** | ***25%*** | ***50%*** | ***75%*** | ***100%*** | ***% of variation explained by first PC*** |
| --- | --- | --- | --- | --- | --- | --- |
| **PC-based composites** | | | | | | |
| Edges | 0.040 | 0.385 | 0.527 | 0.688 | 0.956 | 59.7 |
| Nodes | 0.658 | 0.746 | 0.812 | 0.872 | 0.986 | 83.5 |

#### ***Table S6.* Correlations between edge- and node-based composite scores within NOIs.**

| ***Network*** | ***r***  **PC-weighted Edges & Nodes** |
| --- | --- |
| Global | .108 (.013) |
| Central Executive | .080 (.065) |
| Cingulo-Opercular | .143 (.001) |
| Default Mode | .017 (.695) |
| Hippocampal-Diencephalic | .040 (.360) |
| Multiple Demand | .083 (.056) |
| PFIT | .104 (.017) |
| Salience | .107 (.014) |
| Sensorimotor | .119 (.006) |
| Temporo-Amygdala-Orbitofrontal | .020 (.642) |
| Null | .045 (.298) |

*Note*. P-values are in parentheses.

***Table S7.* Interquartile range for associations between permuted composite scores and cognitive function.** Lower and upper 2.5% boundaries of permuted distributions for associations between permuted composite scores and cognitive function. 95% IQR represents the difference between the upper and lower bounds of each distribution.

| ***Visuospatial Ability*** | | | |
| --- | --- | --- | --- |
| *Element* | *Lower 2.5%* | *Upper 2.5%* | *95% IQR* |
| **Age-based composites** | | | |
| Edges | -0.073 | -0.020 | 0.052 |
| Nodes | -0.391 | -0.374 | 0.017 |
| **PC-based composites** | | | |
| Edges | 0.038 | 0.059 | 0.021 |
| Nodes | 0.380 | 0.387 | 0.007 |
| ***Processing Speed*** | | | |
| *Element* | *Lower 2.5%* | *Upper 2.5%* | *95% IQR* |
| **Age-based composites** | | | |
| Edges | -0.185 | -0.131 | 0.053 |
| Nodes | -0.247 | -0.225 | 0.022 |
| **PC-based composites** | | | |
| Edges | 0.150 | 0.173 | 0.023 |
| Nodes | 0.232 | 0.242 | 0.010 |
| ***Memory*** | | | |
| *Element* | *Lower 2.5%* | *Upper 2.5%* | *95% IQR* |
| **Age-based composites** | | | |
| Edges | -0.070 | -0.015 | 0.055 |
| Nodes | -0.128 | -0.110 | 0.017 |
| **PC-based composites** | | | |
| Edges | 0.032 | 0.054 | 0.022 |
| Nodes | 0.116 | 0.123 | 0.008 |

***Table S8.* Model fit and descriptive statistics for cognitive tests in LBC1936.** Descriptive statistics, intercorrelations, and factor parameters for the cognitive tasks in LBC1936.

| ***Cognitive Task*** | ***Mean*** | ***SD*** | ***Range*** | ***Factor Loading*** | ***Intercorrelations*** | | | | | | | |  |  |  |
| --- | --- | --- | --- | --- | --- | --- | --- | --- | --- | --- | --- | --- | --- | --- | --- |
| **Visuospatial Reasoning** CFI = 0.913; TLI = 0.739; RMSEA = 0.180; SRMR = 0.049 | | | | | **MR** | **BD** | **SSF** | **SSB** | **DSS** | **SS** | **CRT** | **IT** | **LM** | **VPA** | **DB** |
| Matrix Reasoning (MR) | 13.4 | 4.9 | 3 - 25 | 0.663 | 1 |  |  |  |  |  |  |  |  |  |  |
| Block Design (BD) | 34.3 | 10.1 | 10 - 65 | 0.764 | .541 | 1 |  |  |  |  |  |  |  |  |  |
| Spatial Span (Forward) (SSF) | 7.6 | 1.6 | 3 - 12 | 0.462 | .272 | .308 | 1 |  |  |  |  |  |  |  |  |
| Spatial Span (Backward) (SSB) | 7.1 | 1.6 | 1 - 11 | 0.543 | .306 | .398 | .413 | 1 |  |  |  |  |  |  |  |
| **Processing Speed** CFI = 1.0; TLI = 0.999; RMSEA = 0.009; SRMR = 0.011 | | | | |  | | | | | | | | | | |
| Digit-Symbol Substitution (DSS) | 56.6 | 11.6 | 26 - 94 | 0.821 | .333 | .438 | .272 | .271 | 1 |  |  |  |  |  |  |
| Symbol Search (SS) | 25.8 | 6.0 | 4 - 43 | 0.748 | .332 | .467 | .288 | .325 | .620 | 1 |  |  |  |  |  |
| Choice Reaction Time* (CRT) | -64.4 | 8.5 | -108 - -45.9 | 0.645 | .184 | .279 | .276 | .252 | .528 | .473 | 1 |  |  |  |  |
| Inspection Time (IT) | 111.4 | 10.9 | 67 - 134 | 0.462 | .218 | .259 | .218 | .258 | .366 | .341 | .335 | 1 |  |  |  |
| **Memory**  CFI = 1.0; TLI = 1.0; RMSEA = 0.0; SRMR = 0.0 | | | | |  | | | | | | | | | | |
| Logical Memory (LM) | 74.7 | 17.9 | 17 - 116 | 0.784 | .340 | .259 | .160 | .155 | .339 | .279 | .223 | .184 | 1 |  |  |
| Verbal Paired Associates (VPA) | 27.5 | 9.5 | 0 - 40 | 0.684 | .333 | .284 | .157 | .161 | .335 | .259 | .216 | .231 | .530 | 1 |  |
| Digits Backwards (DB) | 7.91 | 2.3 | 2 - 14 | 0.414 | .343 | .311 | .268 | .286 | .354 | .316 | .222 | .175 | .325 | .282 | 1 |

*Note*. Between 1 and 13 participants had missing data across all tests. CRT scores were multiplied by -100 such that higher scores are associated with faster performance. The model for Memory was fully saturated and thus returned perfect model fit.

#### ***Table S9.* UKB-derived age- and PC-weights for connectome elements.**

**Note**: Due to the size of this table, it is only accessible as an Excel file.
